# Adaptation of the *Spalax galili* transcriptome to life under hypoxia may hold a key to a complex phenotype including longevity and cancer resistance

**DOI:** 10.1101/2023.08.01.551427

**Authors:** G. Poetzsch, L. Jelacic, A. Bicker, M. Balling, L. Hellmann, L. Dammer, M.A. Andrade-Navarro, I. Shams, A. Avivi, T. Hankeln

## Abstract

The muroid rodent *Nannospalax galili* (syn. *Spalax*) is adapted to life in underground burrows and tolerates acute exposure to severe hypoxia. Adaptation to hypoxia is correlated with delayed onset of ageing and resistance against tumour formation. *Spalax* becomes five to seven times older than its relatives, the mouse and rat, without displaying signs of ageing or developing ageing-related disorders like cancer. Investigating and understanding adapted genes and gene regulatory networks of *Spalax* might pinpoint novel strategies to maintain an extended healthy phenotype in humans. Here we analysed and compared RNA-Seq data of liver, kidney and spleen of *Spalax* and rat subjected to 6% O_2_ or normoxia. We identified differentially expressed genes and pathways common to multiple organs in *Spalax* and rat. Body-wide differences between *Spalax* and rat affected biological processes like cell death, defence against reactive oxygen species (ROS), DNA repair, energy metabolism, immune response and angiogenesis, which altogether might play a crucial role in *Spalax*’s adaptation to life under oxygen deprivation. In all organs, mRNA expression of genes associated with genome stability maintenance and DNA repair was elevated in *Spalax* compared to rat, accompanied by a lower gene expression of genes associated with aerobic energy metabolism and proinflammatory processes. These transcriptomic changes might be accountable for the extraordinary lifespan of *Spalax* and its cancer resistance. Our results reveal gene regulatory networks that become candidates for the investigation of the molecular bases that underlie the complex phenotype of *Spalax*.

## Introduction

The search for- and selection of non-model organisms with unique adaptive traits harbors the potential to increase our biological insight significantly. In this respect, fossorial rodents, which are adapted to cope with the multiple stresses of underground life, have emerged as attractive systems to study important biomedical conditions related to tissue oxygenation, tumour formation and longevity (Buffenstein et al., 2022) (Lagunas-Rangel, 2018). One of those emerging model taxa is *Nannospalax galili* (also called *Spalax* or *blind mole rat, BMR*), a member of the superspecies *Spalax ehrenbergi* from the muroid family of *Spalacidae*, a group of fossorial rodents that diverged from the phylogenetic lineages leading to mouse and rat approximately 47 million years ago (Fang et al., 2014) (Steppan et al., 2004). The rat-sized *Spalax* individuals inhabit self-dug underground burrows to avoid predators and spend most of their lives solitarily (Lacey et al., 2001). They meet the challenges of their subterranean habitat through multiple evolutionary adaptations. Morphologically, these include a fusiform body shape with a massive spade-formed head for digging (Topachewski, 1976), degenerated subcutaneous eyes which only detect light-dark-differences, a hypertrophied Harderian gland to perceive photoperiodicity (Sanyal et al., 1990) (Pevet et al., 1984) and a sophisticated olfactory system (Burda et al., 1990) combined with an enlarged somatosensory cortex to process tactile stimuli and vibrations for seismic communication (Nevo, 1991). During their speciation in the Pleistocene, all *S. ehrenbergi* subspecies have adapted to aridity and halophyte food resources. In *N. galili*, however, adaptations in kidney weight and urine osmotic concentration are lowly manifested, probably due to its humid habitat in the *Galilean* mountains (Nevo et al., 1989). Due to frequent flooding of its burrow system during seasonal rainfalls in this region, *N. galili* has adapted to overcome the most severe fluctuations in oxygen and carbon dioxide among all *S. ehrenbergi* subspecies. O_2_ concentrations of only 7 % were measured in flooded *Spalax* burrows (Shams et al., 2005). Under laboratory conditions, *N. galili* was capable of surviving 14 h at 3% O_2_, whereas rats died after 2-4 h (Shams et al., 2004). This efficient hypoxia resistance is realized in *Spalax* by a combination of physiological adaptations. As a shared feature of many subterranean rodents, *Spalax* displays a very low basal metabolic rate even under normoxic conditions which leads to a reduction of oxygen consumption (Lacey et al., 2001). In addition, *Spalax* possesses a significantly smaller total skeletal muscle mass than comparatively sized rodents (Widmer et al., 1997). Oxygen distribution, on the other hand, is increased by elevated capillary and mitochondrial density (Avivi et al., 1999). In contrast to *S. ehrenbergi* subspecies living in arid, well-oxygenated and permeable soil, *N. galili* shows elevated concentrations of red blood cells and hemoglobin (Arieli et al., 1986). Together, these features are thought to guarantee unimpaired body functions even under challenging atmospheric oxygen conditions.

Its remarkable hypoxia adaptation is accompanied in *Spalax* by two other striking phenotypic features: Extraordinary longevity and resistance against tumour formation. While laboratory rats have an average lifespan of 3 years (Sengupta, 2013), *Spalax* reaches up to 21 years without displaying clear signs of ageing or age-related disorders (Manov et al., 2013a). Moreover, in 50 years of *Spalax* research, there has never been a reported case of spontaneous tumour formation among thousands of observed individuals. Even exposition to potent carcinogens failed—with very rare exceptions—to induce tumour growth in *Spalax* individuals, whereas 100% of exposed mice and rats in the same study developed tumours (Manov et al., 2013). Interestingly, the blind mole rat shares these important traits with another fossorial rodent, the African naked mole rat *Heterocephalus glaber* (Buffenstein et al., 2022) (Shepard & Kissil, 2020) which is adapted to tolerate moderate-, chronic hypoxia in its populated, eusocial underground colonies, and which reaches lifespans of up to 30 years and develops cancer only in rare singular cases (Buffenstein et al., 2022). Since *Spalax* and *H. glaber* belong to different clades of the rodent phylogeny, they probably evolved these traits convergently during speciation (Steppan et al., 2004) (Fang et al., 2014). It is under current debate that the three combined phenotypes (hypoxia tolerance, longevity, tumour resistance) could be mechanistically linked at the genetic level (Mehta et al., 2009) (Boretto et al., 2018) (Rascón & Harrison, 2010) (Snell & Johnston, 2014) (Strzyz, 2023). Comparative genomics and molecular analyses in recent years have provided evidence that both positively selected sequence changes in coding genes and mutations in gene regulatory elements have contributed to adaptation in the subterranean rodents (Gladyshev et al., 2011) (Fang et al., 2014) (Malik et al., 2016) (Schmidt et al., 2017) (Davies et al., 2018). In *Spalax*, for example, fixed amino acid replacements in the crucial tumour suppressor p53 render the protein unable to activate some of its apoptosis-regulating target genes (Ashur-Fabian et al., 2004). Additionally, *Spalax* p53 over-activates genes associated with cell-cycle arrest. This suggests an adaptive strategy to prevent hypoxia-induced cell death, while allowing DNA repair to take place. Pseudogenization, e.g., associated with visual perception and DNA repair, is another evolutive feature of *Spalax* observed at the genomic level (Zheng et al., 2022). Additional gene-focused studies revealed substantially increased mRNA and protein expression of key players involved in a variety of functions in *Spalax* organs in comparison to tissues from the non-hypoxia-adapted rat, e.g., elevated Hif1 (Shams, Nevo, et al., 2005) and Epo levels (Shams et al., 2004) or overexpression of oxygen supply proteins (Avivi et al., 2010), or ROS defence enzymes (Schülke et al., 2012) and tissue vascularization genes (Vegf, (Avivi et al., 2005)). Large-scale transcriptome analyses of a subset of *Spalax* organs uncovered additional gene-regulatory adaptations, e.g., suggesting an attenuated transcriptional response of energy metabolism genes under normoxic conditions in brain (Malik et al., 2016) and in response to hypoxia in liver tissue (Schmidt et al., 2017). Moreover, *Spalax* liver and brain transcriptomes (in comparison to rat) revealed constitutively increased expression of DNA damage repair genes, including important players such as the base excision repair-associated *glycosylase gene Endonuclease VIII-like 2* (*Neil2*), the *Xeroderma pigmentosum-associated* (*Xpa*) nucleotide excision repair gene and the progeria-associated helicase Werner (*Wrn*) (Malik et al., 2016) (Schmidt et al., 2017). In addition, replicative stress response genes known to interact with *Wrn*, such as the *Ataxia telangiectasia and Rad3 related gene* (*Atr*), the *Serine/Threonine Kinase Ataxia Telangiectasia Mutated gene* (*Atm*) and members of the *Fanconi-Anemia pathway*, showed increased mRNA expression in *Spalax* livers, compared to rat (Schmidt et al., 2017). Altogether, the observed genetic and gene regulatory changes may facilitate genome stability in *Spalax* under conditions where severe hypoxia and subsequent reoxygenation challenge tissues and cells, e.g., via the production of toxic reactive oxygen species (Malik et al., 2016).

The results above support that increased genome stability explains decreased ageing and tumourigenesis as “side effects” of hypoxia adaption in *Spalax*. We hypothesized that given the abundant phenotypic and molecular evidence of the adaptation of *Spalax* and other fossorial species living in similar conditions to hypoxia described above, additional evidence should be found across other tissues such as kidney and spleen, which collectively could improve our understanding of the general molecular principles of the relation between hypoxia, ageing and cancer. To investigate this, we generated and interpreted RNA-Seq data from additional *Spalax* and rat organs (kidney, spleen) from animals subjected/exposed to hypoxia (20 % O_2_ > 6 % O_2_) and normoxia controls. Including our previous liver RNA-seq data, we then conducted meta-transcriptomic comparisons across *Spalax* tissues to elucidate overarching versus tissue-specific gene regulatory responses. We thereby aimed at inferring common, body-wide differences in gene expression, which might have played a crucial role in adaptation to life under hypoxic stress conditions. We discuss the relevance of our results for the elucidation of the molecular mechanisms of ageing and cancer.

## Materials and Methods

### Sample preparation

Six *N. galili* specimens were captured from basalt, heavy soil in fields of northern Israel and kept in the animal house of the *Institute of Evolution, Haifa University* in accordance with the regulations of the Israel *Nature and Park Authority, Science and Conservation Unit*. To investigate the impact of O_2_ deprivation on the transcriptomes of different organs in *Spalax* and rat, three *Spalax* (female) and three rat individuals (male) were exposed to 6 % O_2_ for six hours in 70x70x50 cm chambers divided into separate cells at a gas flow rate of 3.5 l/min. None of the animals died during the hypoxia exposure. Afterwards, they were sacrificed immediately by Ketaset CIII (Fort Dodge, USA) injection. From each individual, kidney and spleen were dissected. Liver samples from the same animals were analysed prior to this study (Schmidt et al., 2017). All animal experiments were approved by the University of Haifa Ethics Committee (Permit #193/10). RNA was extracted from kidney and spleen with the RNeasy lipid tissue kit (Qiagen, Hilden, Germany), including the RNase-Free DNase I treatment (Qiagen). RNA integrity was validated on a 2100 Bioanalyser with the Agilent RNA 6000 Nano kit (Agilent Technologies, Santa Clara, USA). Sequencing libraries were generated with the Illumina TruSeq RNA library Prep kit v2 (StarSEQ, Mainz, Germany). All libraries were sequenced as 100 bp paired-end reads on an Illumina HiSeq2500 platform, run by the NGS core facility of the Biology Department of Johannes Gutenberg University Mainz (Germany). Sequence data were uploaded for public availability to ENA (Acc. No. PRJEB64658).

### RNA-Seq analyses

RNA-Seq data were quality-trimmed and filtered for adapter sequences with Trimmomatic v.0.3.6 (Bolger et al., 2014) using a sliding window approach with a window size of 4 nt and a quality score cutoff of 20. The first 13 nt of each read were cropped and reads shorter than 20 bp discarded. The processed reads were mapped with the STAR aligner v.2.5.3 (Dobin & Gingeras, 2016)) to the respective annotated reference genomes of *Rattus norvegicus* (6.0.92; http://ftp.ensembl.org/pub/current_fasta/rattus_norvegicus/dna/) and *N. galili* (1.0.92; http://ftp.ensembl.org/pub/current_fasta/nannospalax_galili/dna/). Count data files and index files were generated, using a sjdbOverhang of 87. The mismatch maximum was set to 0.04 per base and a multimap filter was applied with a maximum of 20 matches.

Gene IDs of *Spalax* and rat were matched by identifying respective orthologues for Ensemble IDs with *BLASTp*. A BLOSUM45 matrix was chosen, at a 100 amino acid cutoff and a minimum identity of 30 %. Only those genes that matched the top BLAST hits as judged by the E-value for both search directions, i.e., *Spalax* query searches in rat databases and *vice versa*, were used.

Differential gene expression was determined between samples with DeSeq2 v.1.30.1 (Love et al., 2014). A padj. cutoff of < 0.05 was used to define differential gene expression and a Log2 FoldChange cutoff of greater than zero (> 0) for up- and lower than zero (< 0) for down regulation. For interspecies comparison, counts were normalized with regard to the exon sum length using the *normMatrix* function as part of the DESeq2 package. A previously published RNA-Seq dataset (Schmidt et al., 2017) was re-evaluated using the same analytical tools to compare the liver transcriptome. Potential biases in gene expression (sex, species, strain, age) were accounted for in the previous study by Schmidt et al. (2017) via comparison with public rat, mouse and human RNA-Seq data. Based on these comparisons a substantial effect of such biases on interspecies- and hypoxia-related gene expression changes is improbable. The differential expression data was mapped onto relevant *KEGG pathways* for visualization using the *pathview* R package (Luo & Brouwer, 2013). Principal component analyses were generated with in R, using the function plotPCA. Resulting gene lists were compared between species and between different O_2_-conditions using the R VennDiagram package (Chen & Boutros, 2011). We validated the presence of expected organ-specific transcription patterns by comparison with annotated genes that display tissue-specific overexpression(Yu et al., 2014). RNA-Seq data were analysed with the use of QIAGEN Ingenuity Pathway Analysis (QIAGEN Inc., https://digitalinsights.qiagen.com/IPA; (Krämer et al., 2014)). A padj. cutoff of p < 0.05 was used to define pathway enrichment and a z-score cutoff of < -1 or > 1 for inactivation and activation, respectively.

### Validation of RNA-Seq data by qRT-PCR

Differential expression of a subset of candidate genes was validated by quantitative real-time reverse-transcriptase PCR (qRT-PCR). 600 ng of quality-checked RNA of all samples was used to synthesize cDNA with the Superscript III RT kit (Thermo Fisher, Waltham, USA). To generate a standard-curve for determining absolute RNA copy numbers of each tested gene, plasmids containing corresponding amplicons were generated: PCRs were conducted in a peqSTAR 96 Thermocycler (Peqlab, Erlangen, Germany) with the TrueStart Taq DNA Polymerase kit (Thermo Fisher, Waltham, USA) and purified with the Wizard SV Gel and PCR clean up System (Promega, Madison, USA). PCR products were cloned into pGem-T Easy vectors (Promega), transformed in *E. coli* DH10B cells, plated out and purified with the *GeneJET Plasmid Miniprep Kit* (Thermo Fisher). To validate the correct inserts, plasmids were Sanger-sequenced (StarSEQ, Mainz, Germany). qRT-PCR was conducted with the GoTaq qPCR Master Mix (Promega) at a total volume of 10 µl in an ABI 7500 Fast Real Time PCR cycler (Applied Biosystems). For absolute quantification, cDNAs of all samples, measured in triplicate, were normalized to plasmid-copies, which were measured in serial 10-fold dilutions.

## Results

### Transcriptome data metrics

To unveil the potential adaptive mechanisms which allow the long-lived rodent *N. galili* to tolerate severe hypoxia on the gene regulatory level, we subjected n = 3 individuals of both species to either 6 % O_2_ or to normoxia (room air) for six hours and dissected liver, kidney and spleen tissues. RNA was isolated to generate transcriptome-wide mRNA expression profiles by Illumina sequencing. Between 10 and 54 million reads per dataset were mapped to their respective reference genomes. (Tab. S1). Expected organ-specific gene expression patterns as described by Yu et al. (2014) were clearly observable within our data (S2).

We detected expression in at least one of the biological replicates for at least 16350 genes in *Spalax* and at least 17700 genes in rat in all analysed tissues under both oxygen conditions (S3). When comparing the gene expression for each tissue via Principal Component Analyses (PCA), we observed that replicate samples (n = 3) of the same species subjected to the same oxygen conditions clustered together as expected (Fig. S4). In all tissue comparisons, the interspecies variance between *Spalax* and rat was larger than between the hypoxia- and normoxia-treated samples of the same species, implying that hypoxia induction had less impact on the RNA expression profile than species divergence.

### Quantitative transcriptome differences in the hypoxia response of *Spalax* and rat

To study the magnitude of the gene regulatory response after hypoxic stress, we identified differentially expressed genes (DEGs) between normoxia and hypoxia for all three tissues in both species (padj. < 0.05). In all tissues, the hypoxia response was notably stronger in rat than in *Spalax* (Fig. 1A). After hypoxia treatment, about 1.5 times more genes were dysregulated in rat liver, 1.45 times more in rat kidney and 1.6 times more in rat spleen, as compared to *Spalax*. Fig. 1B shows tissue-wise scatterplots of the hypoxia induction of each gene for rat and *Spalax*. In particular in spleen, many more dysregulated genes (red dots) were observed in rat (3326) compared to *Spalax* (2055). Together, these data indicate a stronger response to hypoxic stress of rat tissues than of the hypoxia-adapted *Spalax*.

**Fig. 1.**
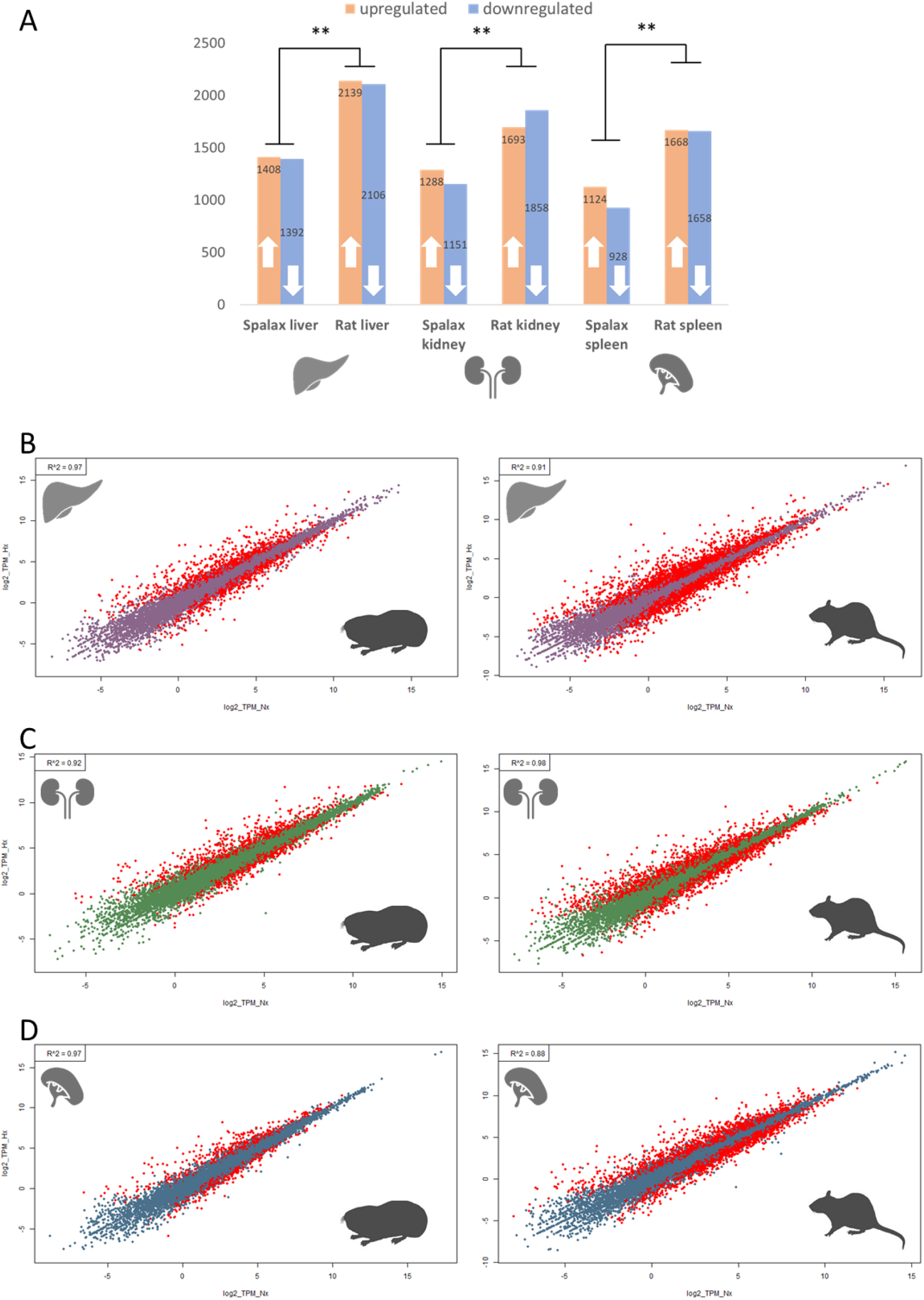
(A): Number of Spalax and rat DEGs after hypoxic stress in liver, kidney and spleen. (B-D): Scatterplot visualization of DEGs following hypoxic treatment in Spalax and rat liver, kidney and spleen. In each graph, log 2 TPM fold-changes are plotted. Differentially expressed genes (padj <0.05) are marked in red.

In addition to the observed quantitative differences, the sets of genes induced by hypoxia differed qualitatively between the two species. From those genes differentially expressed under hypoxia in *Spalax*, only 46 % (liver), 33 % (kidney) and 37 % (spleen) were also regulated in the rat (Fig. S5). This suggests that both organisms transcribe a large set of species-specific hypoxia-responding genes. Ca. 32 % (1677) and 39 % (2897) of hypoxia-stimulated DEGs were common to at least two organs in *Spalax* and rat, respectively (Fig. 2). These genes might represent a systemic transcriptional response towards hypoxic stress throughout different tissues.

**Fig. 2:**
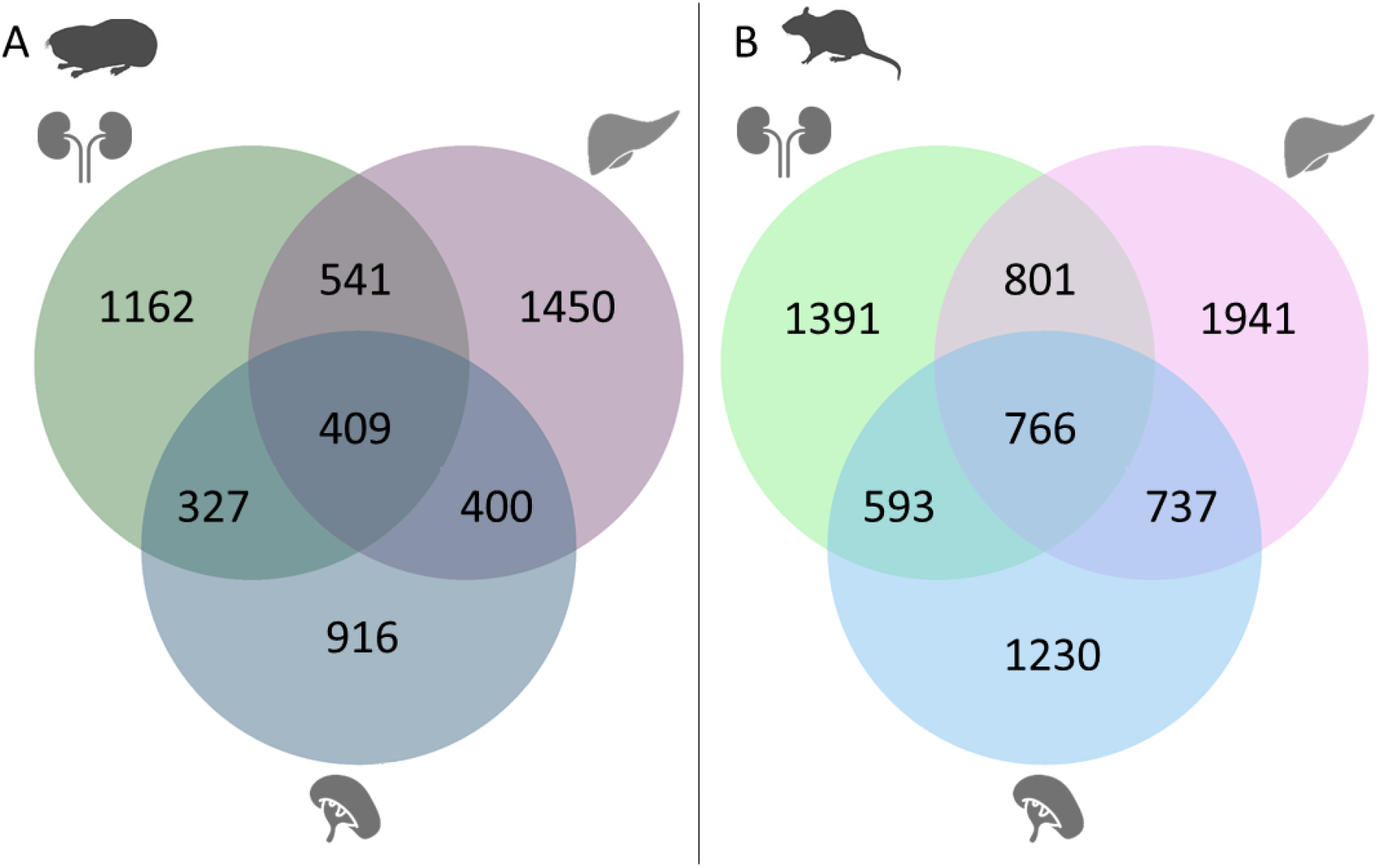
Venn diagram of hypoxia-regulated genes compared between all three organs (padj < 0.05). A: Genes differentially expressed under hypoxia in Spalax; B: Genes differentially expressed under hypoxia in rat

#### Functional classification of genes involved in the hypoxia response

To interpret the biological function of the DEGs after hypoxia exposure in *Spalax* and rat, pathway enrichment analyses were performed for each of the three organs (S6). Identified pathways describing biological processes were summarized for comparison. Besides an enrichment of general cellular signaling processes in both species, pathway enrichment results of **liver** (Tab. 1) tissue showed that in both species hypoxia-activated pathways were associated with cell death (e.g. “Senescence Pathway”), proinflammatory processes (e.g. “Acute Phase Response Signaling”), hypoxia stress response (e.g. “HIF1α Signaling”), extracellular matrix (ECM) and angiogenesis (e.g. “Tumour Microenvironment Pathway”). In rat, but not in *Spalax*, enrichment in metabolism associated terms like “Gluconeogenesis I” and “Glycolysis I” was detected in hypoxia activated genes.

**Table 1:**
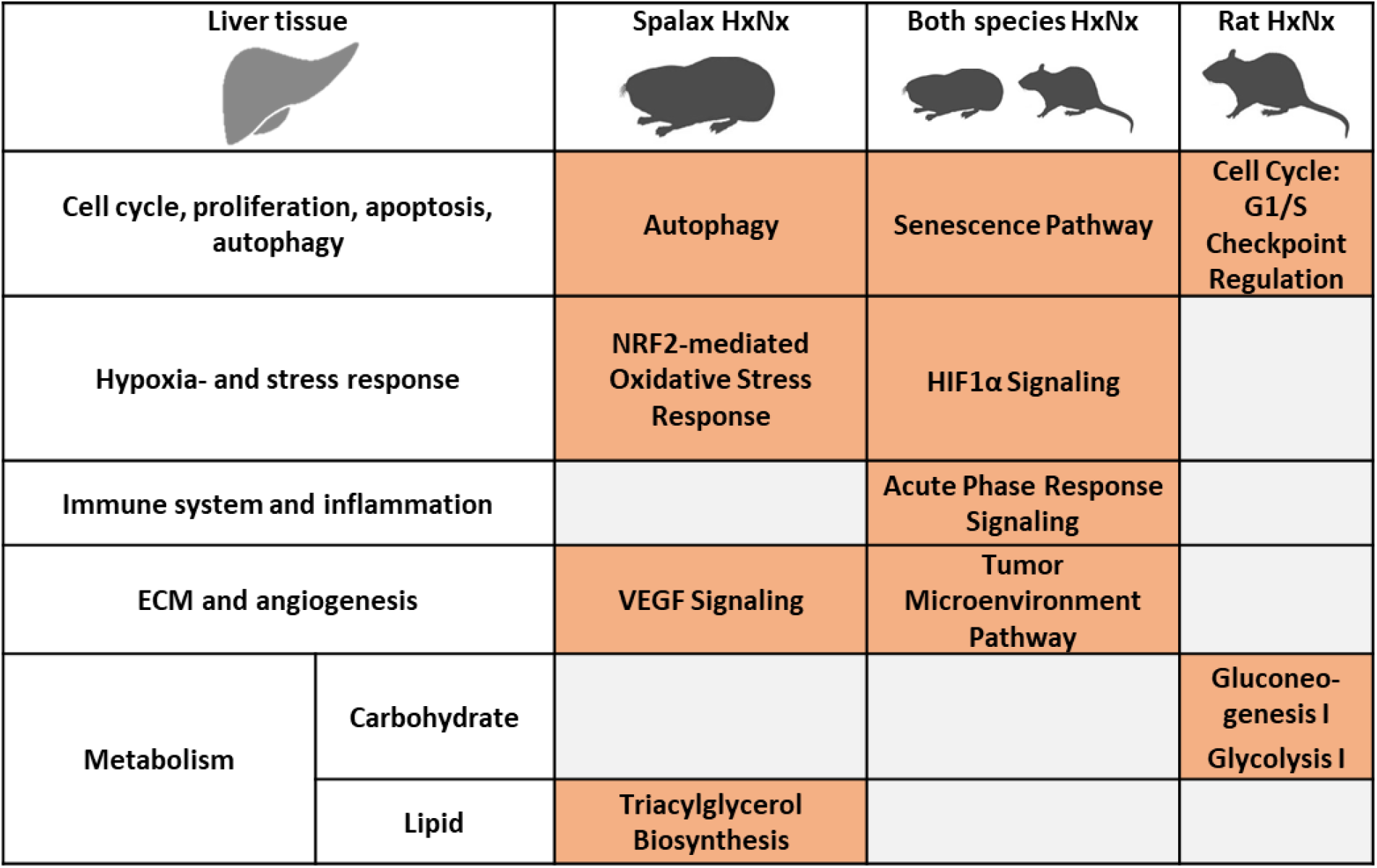
Selected pathways induced by hypoxia in the liver of Spalax and rat (padj < 0.05). Pathway enrichment was predicted based of the differential expression between hypoxic and normoxic conditions (HxNx) in the respective species. Pathways are grouped into larger categories (left margin) for improved overview (orange = predicted activation (z-score > 1)

| Liver tissue |  | Spalax HxNx | Both species HxNx | Rat HxNx |
| --- | --- | --- | --- | --- |
| Cell cycle, proliferation, apoptosis, autophagy |  | Autophagy | Senescence Pathway | Cell Cycle: G1/S Checkpoint Regulation |
| Hypoxia- and stress response | | NRF2-mediated Oxidative Stress Response | HIF1α Signaling | |
| Immune system and inflammation |  |  | Acute Phase Response Signaling |  |
| ECM and angiogenesis |  | VEGF Signaling | Tumor Microenvironment Pathway |  |
| Metabolism | Carbohydrate |  |  | Gluconeogenesis I<br>Glycolysis I |
|  | Lipid | Triacylglycerol Biosynthesis |  |  |

In **kidney**, in addition to the regulation of general cellular signaling processes, hypoxic activation of the following pathways was observed in both species (Tab. 2): “Wound healing Signaling Pathway”, angiogenesis and ECM formation (”VEGF-Signaling”, “Tumour Microenvironment Pathway”), hypoxia- and oxidative stress-associated processes (“HIF1α Signaling”, “NRF2-mediated Oxidative Stress Response”), proinflammatory pathways (“Acute Phase Response Signaling”) and metabolism-associated terms (e.g. “Type II Diabetes Mellitus Signaling”). Particularly in *Spalax*, the activation of many cell-death associated terms was detected (e.g., “MYC Mediated Apoptosis Signaling).

**Table 2:**
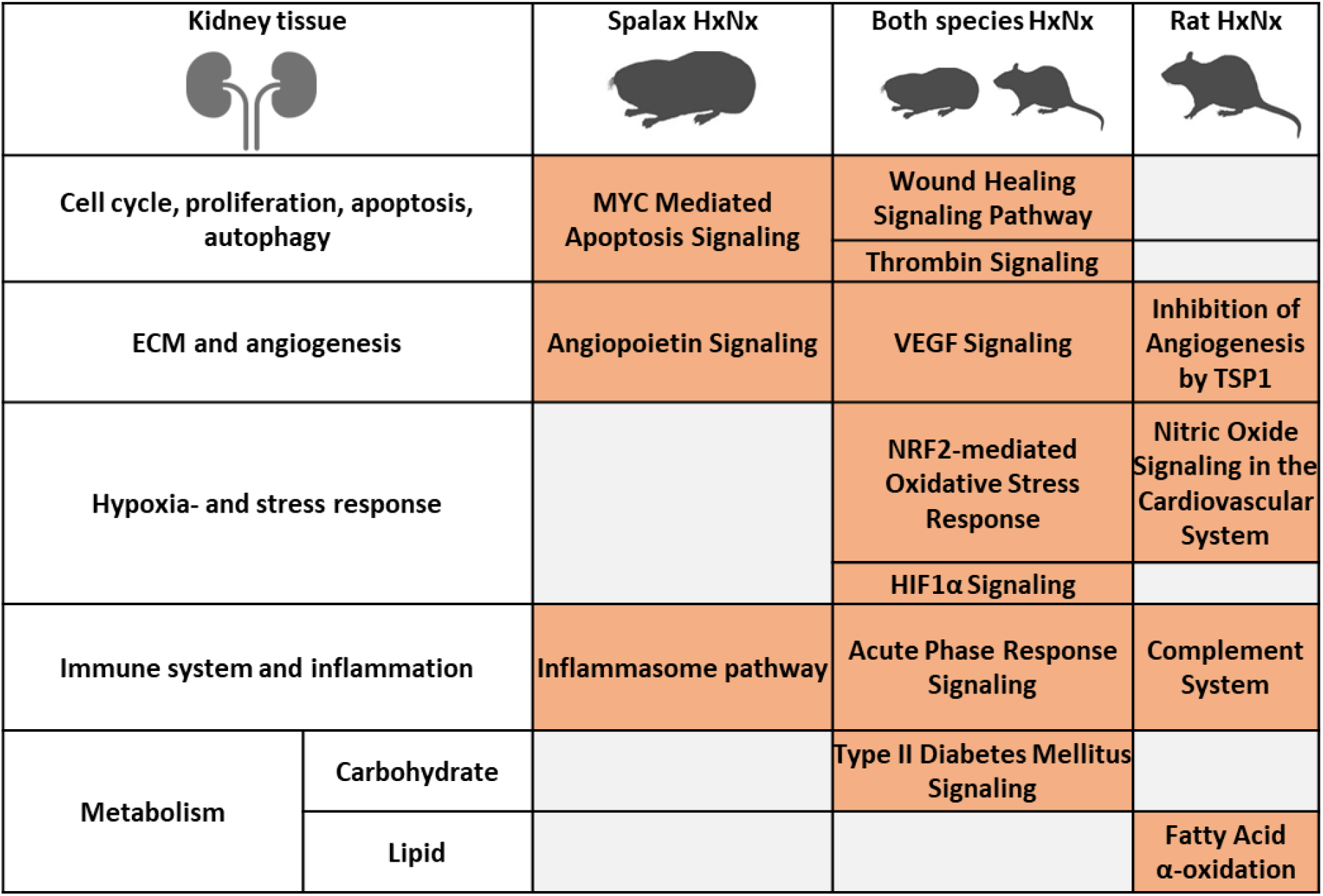
Selected pathways induced by hypoxia in the **kidney** of Spalax and rat (padj < 0.05). Pathway enrichment was predicted based of the differential expression between hypoxic and normoxic conditions (HxNx) in the respective species. Pathways are grouped into larger categories (left margin) for improved overview (orange = predicted activation (z-score > 1)

| Kidney tissue |  | Spalax HxNx | Both species HxNx | Rat HxNx |
| --- | --- | --- | --- | --- |
| Cell cycle, proliferation, apoptosis, autophagy |  | MYC Mediated Apoptosis Signaling | Wound Healing Signaling Pathway |  |
|  |  |  | Thrombin Signaling |  |
| ECM and angiogenesis |  | Angiopoietin Signaling | VEGF Signaling | Inhibition of Angiogenesis by TSP1 |
| Hypoxia- and stress response |  |  | NRF2-mediated Oxidative Stress Response | Nitric Oxide Signaling in the Cardiovascular System |
| | | | HIF1α Signaling | |
| Immune system and inflammation |  | Inflammasome pathway | Acute Phase Response Signaling | Complement System |
| Metabolism | Carbohydrate |  | Type II Diabetes Mellitus Signaling |  |
| | Lipid | | | Fatty Acid α-oxidation |

Pathway enrichment results of **spleen** tissue (Tab. 3) indicated that in both species hypoxia-responsive DEGs were enriched in terms related to the activation of signaling pathways that are associated with pathologies and damage (“Wound Healing Signaling Pathway”, “Osteoarthritis Pathway”, “IL6-Signaling”). In notable contrast to the other organs, hypoxia response pathways (“HIF1α Signaling”,” Nitric Oxide Signaling in the Cardiovascular System”) were specifically observed to be activated in rat and not in *Spalax*.

**Table 3:**
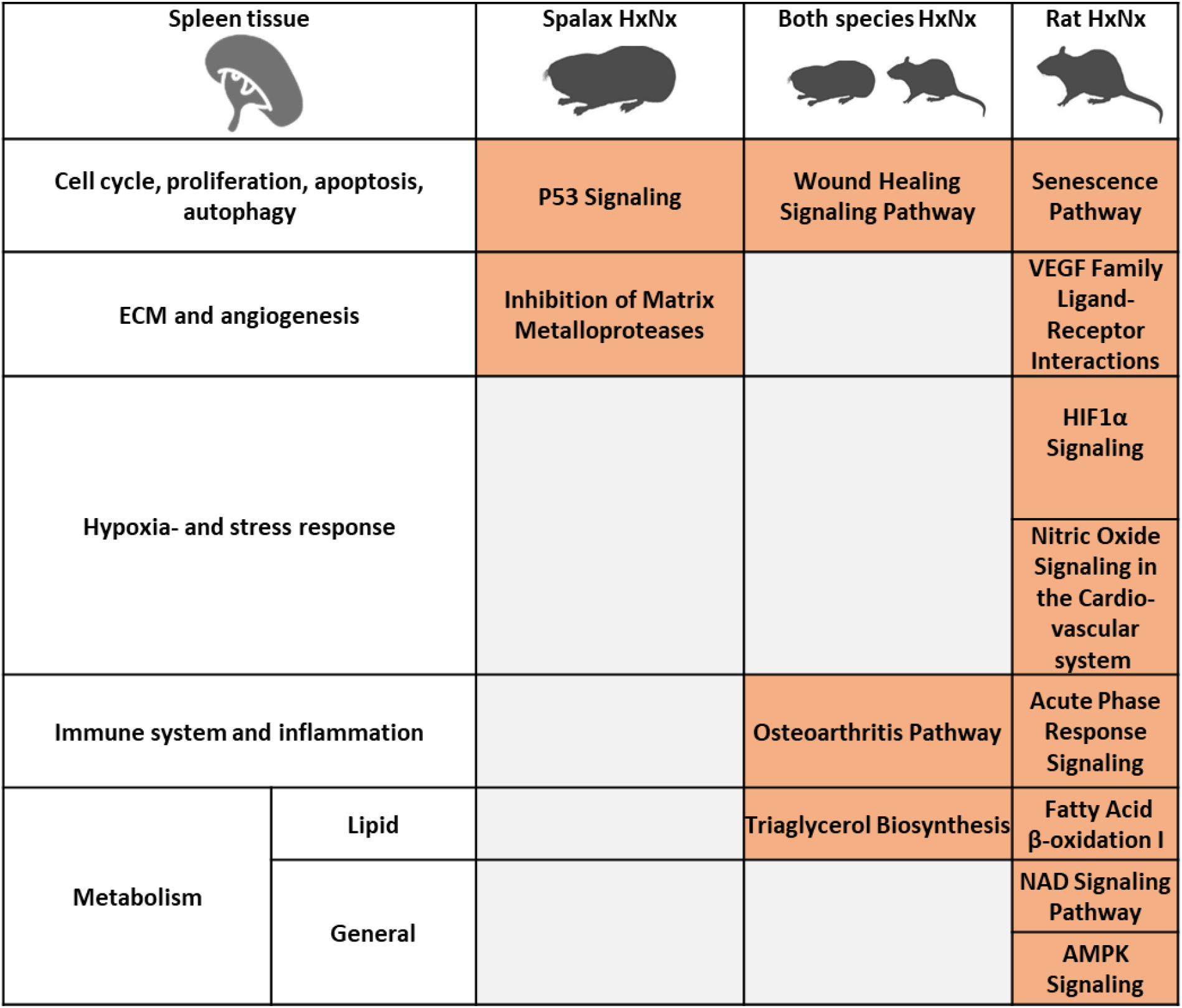
Selected pathways induced by hypoxia in the **spleen** of Spalax and rat (padj < 0.05). Pathway enrichment was predicted based of the differential expression between hypoxic and normoxic conditions (HxNx) in the respective species. Pathways are grouped into larger categories (left margin) for improved overview (orange = predicted activation (z-score > 1)

| Spleen tissue |  | Spalax HxNx | Both species HxNx | Rat HxNx |
| --- | --- | --- | --- | --- |
| Cell cycle, proliferation, apoptosis, autophagy |  | P53 Signaling | Wound Healing Signaling Pathway | Senescence Pathway |
| ECM and angiogenesis |  | Inhibition of Matrix Metalloproteases |  | VEGF Family Ligand-Receptor Interactions |
| Hypoxia- and stress response | | | | HIF1α Signaling |
|  |  |  |  | Nitric Oxide Signaling in the Cardio-vascular system |
| Immune system and inflammation |  |  | Osteoarthritis Pathway | Acute Phase Response Signaling |
| Metabolism | Lipid | | Triaglycerol Biosynthesis | Fatty Acid β-oxidation I |
|  | General |  |  | NAD Signaling Pathway |
|  |  |  |  | AMPK Signaling |

The results indicated that many processes important for hypoxia response are activated in both, the adapted rodent *Spalax* and the non-adapted rat. To determine whether this apparent functional similarity in hypoxia response is orchestrated by the regulation of *the same genes* in *Spalax* and rat, we compared the regulation of targeted genes from hypoxia-responsive pathways like “HIF1α Signaling” or “NRF2-mediated Oxidative Stress Response” between *Spalax* and rat in all analysed organs (S7). In liver and kidney, both species regulated around half of the 139 analysed hypoxia-associated genes, while in spleen 50 genes were hypoxia-responsive in the rat and an even lower number (19) in *Spalax*. Only 9 genes showed a hypoxia-induced regulation in both species and all three organs. In an organ-wise comparison, 28 genes in liver, 21 in kidney and no genes in spleen were regulated in the same direction in both species. On the other hand, 39 genes in liver, 40 in kidney and 42 in spleen were regulated in only one of the two species e.g. *Hypoxia-inducible factor-2alpha* Epas1 which was only upregulated in the liver of rat but not of *Spalax* or the *E3 ubiquitin-protein ligase* Mdm2 which was activated *Spalax*-specifically. Among this set were also genes that showed an opposing direction of regulation in *Spalax* and rat like the *MAP kinase-interacting serine/threonine-protein kinase 1* Mknk1 which was upregulated in the liver of rat in response to hypoxia but downregulated in *Spalax*.

In conclusion, the transcriptome data reveal that around 1.5-fold-more genes are regulated in rat compared to *Spalax* in response to hypoxia. The majority of regulated genes differs between *Spalax* and rat, yet this results in the activation of similar biological pathways in liver and kidney. In case of the hypoxia and stress-response pathways, many genes showed species-specific response patterns. In spleen, the hypoxia-response was attenuated in both species, but especially in *Spalax*. Our results indicate an adapted and potentially more efficient gene regulatory hypoxia-response in *Spalax*.

### Differences in gene expression under normoxic conditions between *Spalax* and rat

Differences between *Spalax* and rat in gene regulation may not only manifest themselves during an acute response to hypoxic stress, but also “constitutively” at normoxic conditions. In all three organs, more than half of the genes were differentially expressed between *Spalax* and rat at normoxia (Fig. S8), the number being evenly distributed between higher- and lower-expressed genes. Similar to the patterns detected in the transcriptional hypoxia response, most genes (74 % of all DEGs, n = 9524) were differentially expressed between the two species in at least two organs. Around 37 % of the DEGs (n = 4781) were differentially expressed in all three organs, indicating global adaptations on the gene regulatory level in *Spalax* compared to rat (Fig. 3).

**Fig. 3:**
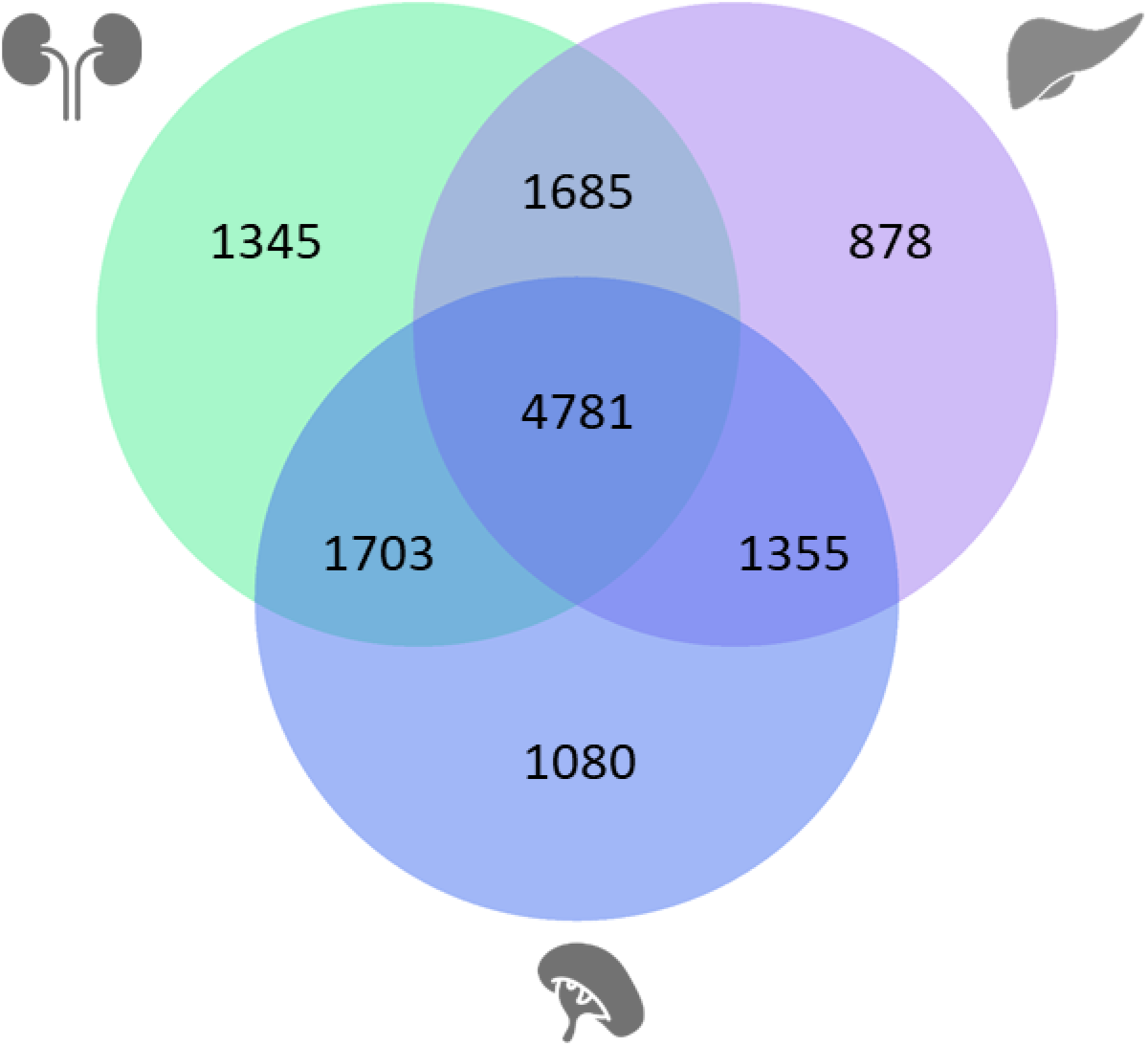
Differential gene expression in Spalax versus rat under normoxic conditions between all three organs (padj < 0.05)

#### Functional classification of genes differentially expressed between Spalax and rat at normoxia

Interspecies DEGs at normoxia were analysed for pathway enrichment to interpret their biological function (S6). We observed a trend of inactivation of processes associated with proliferation e.g., (“mTOR signaling”), proinflammatory pathways (e.g., “Neuroinflammation Signaling Pathway”), coagulation (“Thrombin Signaling”) and extracellular matrix components (e.g., “Dermatan Sulfate Degradation”) and a trend of activation of DNA repair-associated pathways (e.g., “Base excision repair”) in multiple organs in *Spalax* compared to rat (Tab. 4).

**Table 4:**
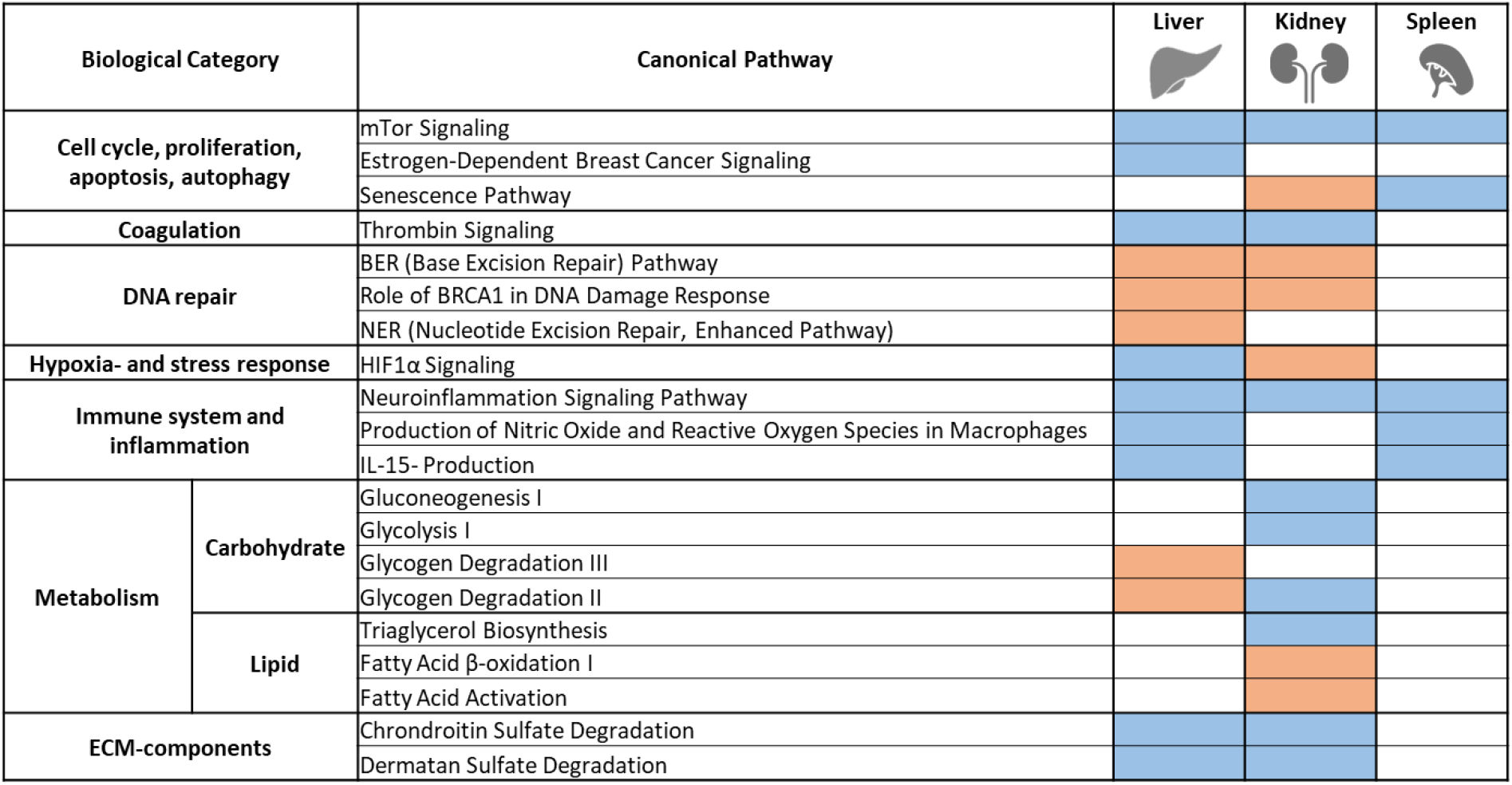
Selected pathways differentially expressed between Spalax and rat (Spalax/rat) (padj < 0.05) in normoxic liver, kidney and spleen. Pathways are grouped into larger categories (left margin) for improved overview (orange = predicted activation (z-score > 1), blue = predicted inactivation (z-score < -1).

Enrichment results in **liver** tissue specifically showed inactivation of hypoxia-associated pathways (“HIF1α Signaling”) and activation of metabolic pathways (“Glycogen Degradation II-“, “Glycogen Degradation III-“).

In **kidney**, “HIF1α Signaling” was predicted to be activated, while carbohydrate metabolism-associated pathways (“Glycolysis”,” Gluconeogenesis”, “Glycogen Degradation”) were predicted to be inactivated. In kidney, we specifically observed enrichment of genes associated with lipid metabolism (“Triacylglycerol Biosynthesis” and “Fatty Acid β-oxidation I”). In the spleen, activity changes for DNA-repair- or other metabolism-associated processes were not observed.

In conclusion, the enrichment analysis of pathways differentially expressed between *Spalax* and rat pinpointed several processes that showed the same activation status across organs. Detailed inspection of such pathways, e.g., mTOR (Tab. 5), in fact, revealed that the same genes were differentially expressed in more than one organ by similar means, i.e., direction of regulation and amplitude, suggesting the action of body-wide gene-regulatory mechanisms.

**Table 5:**
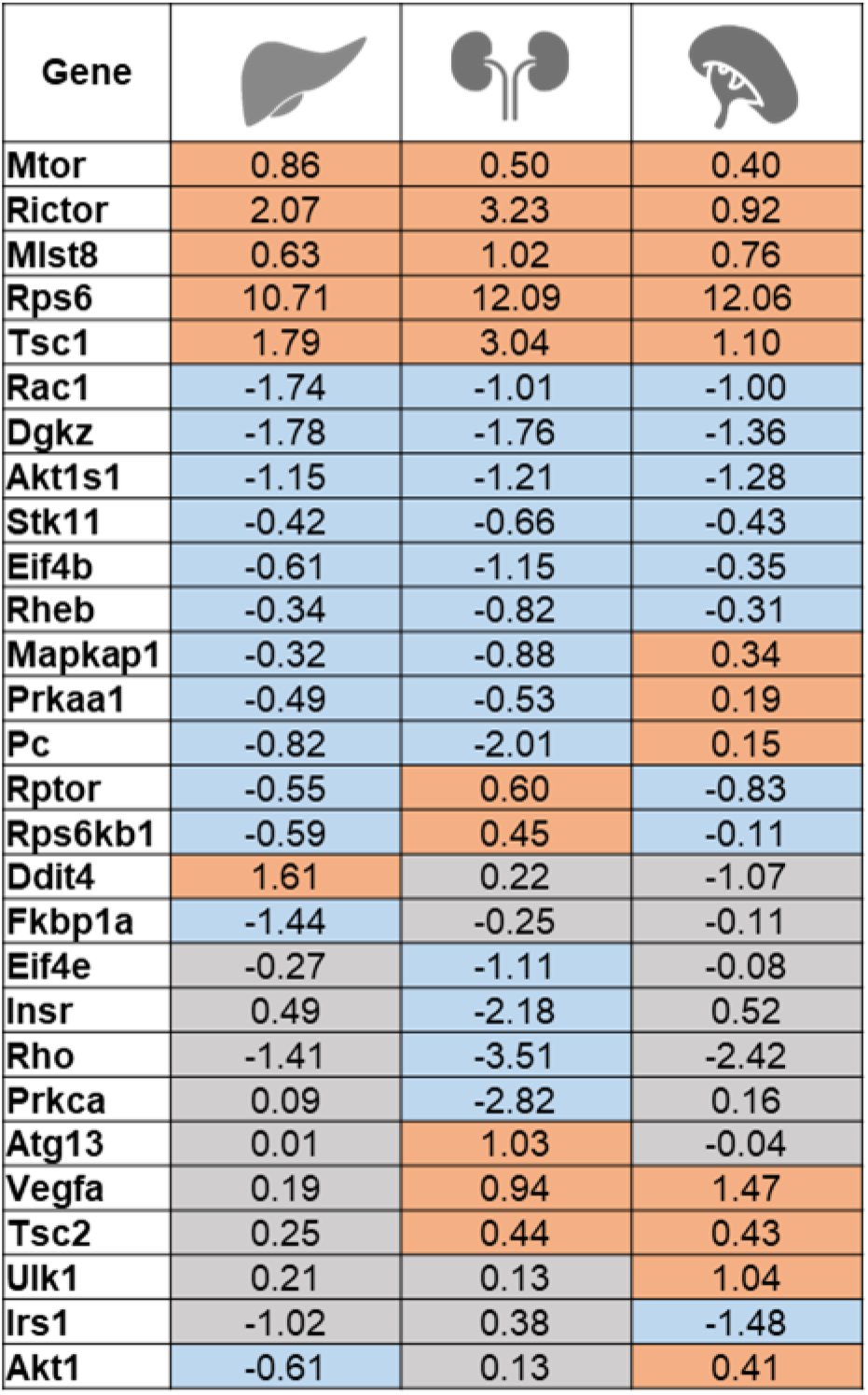
Genes contributing to mTor signaling and their respective fold-change for the differential expression between Spalax and rat under normoxic conditions. Differentially expressed between Spalax and rat in normoxic liver, kidney and spleen. Orange = elevated expression in Spalax (Log_2_FC > 0), blue = lower expression in Spalax (Log_2_FC < 0), grey = differential expression was not significant (padj. > 0.05)

#### Validation of the RNA-Seq analysis via qRT-PCR

Based on the results of the functional enrichment analyses and selection of important candidate genes, we studied the hypoxia inducibility and interspecies expression differences of a selected gene set representing different functional categories (A2m, Atr, Cisd2, Fgf21, Wrn, Xpa, Rcan1, Gpnmb, Fen1, Hmox1, Vegfa, Pnkp) by quantitative realtime RT-PCR (S9). We could confirm the direction of all relevant gene expressional changes in the respective organs.

## Discussion

Subterranean rodents like the blind mole rat *(Nanno)Spalax* spec. exhibit a fascinating combination of phenotypes, linking their adaptation to environmental hypoxia with a remarkable tumour resistance and longevity (Buffenstein et al., 2022) (Lagunas-Rangel, 2018). Previous studies, addressing single genes and the complete genome sequences revealed interesting mutations in candidate coding sequences, but also suggested that differences in gene regulation may underlie adaptations in the subterranean species (MacRae et al., 2015) (Davies et al., 2018) (Schmidt et al., 2017) (Kurz et al., 2017) (Heinze et al., 2018). Here, we generated and compared RNA-Seq datasets from vital organs of the hypoxia-tolerant blind mole rat *S. galili* and –for comparison– from the hypoxia-sensitive rat. We hypothesized that, in particular, organ-overarching patterns of gene expression might yield insights into the molecular pathways that are responsible for the complex phenotype of *Spalax*.

### RNA-Seq data suggest body-wide adaptations at the transcriptome level

Applying a threshold of 2-fold, we measured changes in gene expression in response to hypoxia for 1340/1087/808, and 2126/1678/1504 genes in liver, kidney and spleen of *Spalax* and rat, respectively. These numbers are in line with previous RNA-Seq based transcriptome studies of hypoxia-exposed rodents (Schmidt et al., 2017) (Dong et al., 2020) (Wang et al., 2021). In an interspecies comparison at normoxia, 6426 (liver), 7173 (kidney) and 6205 (spleen) genes showed an expression difference of at least 2-fold between the two species. Comparable studies of the liver transcriptome of the hypoxia-adapted naked mole rat and its non-adapted hypoxia-sensitive relative, the guinea pig, reported differential expression of > 4000 genes between the two taxa (Heinze et al., 2018). Considering differences in the experimental setup and the closer phylogenetic distance between naked mole rat and guinea pig (39.5 mya) compared to *Spalax* and rat (47.4 mya) (Lewis et al., 2016), our results on the interspecies expression differences are also in the expected range. Measurements of RNA level can serve as a reliable proxy for estimating gene expression and can account for approximately two thirds of the observed protein abundance in mammalian cells (Vogel et al., 2010). In the naked mole rat, a similar correlation between the RNA and the protein level in an interspecies comparison has been previously reported (Heinze et al., 2018). However, the same study showed that for some key adaptive processes such as oxidative phosphorylation, differential expression only manifested itself at the protein level. In the future, it will thus be necessary to integrate our transcriptomics approach with protein data to more comprehensively understand adaptation of gene regulation in *Spalax*.

### Complex species-specific patterns characterize transcriptional gene regulation in the rodent models

As a general trend, visible in all three organs studied, a significantly higher number of genes responded to hypoxic stress in the hypoxia-sensitive rat as compared to the hypoxia-tolerant *Spalax* (comp. Fig 1). At the same time, the majority of hypoxia-responsive genes differed between the two rodents (Fig. S5). Still, in many cases, pathways associated with similar biological functions such as response to hypoxia, proliferation control, angiogenesis or immune response were activated in both species (Fig. 4). Detailed investigation of the expression of genes contributing to the respective pathways revealed that often different genes were regulated in the two species within a respective biological pathway, while in other cases the same genes showed opposing directions of regulation in *Spalax* and rat. These results indicate that, while classical hypoxia response pathways are activated in both rodents, the adapted *Spalax* achieves a biological response by utilizing overall less genes. The regulation of different genes compared to rat as exemplified in the mTor signaling pathway, indicates a potentially more sophisticated gene regulatory response which could conserve energy by utilizing less resources for gene expression.

**Fig. 4:**
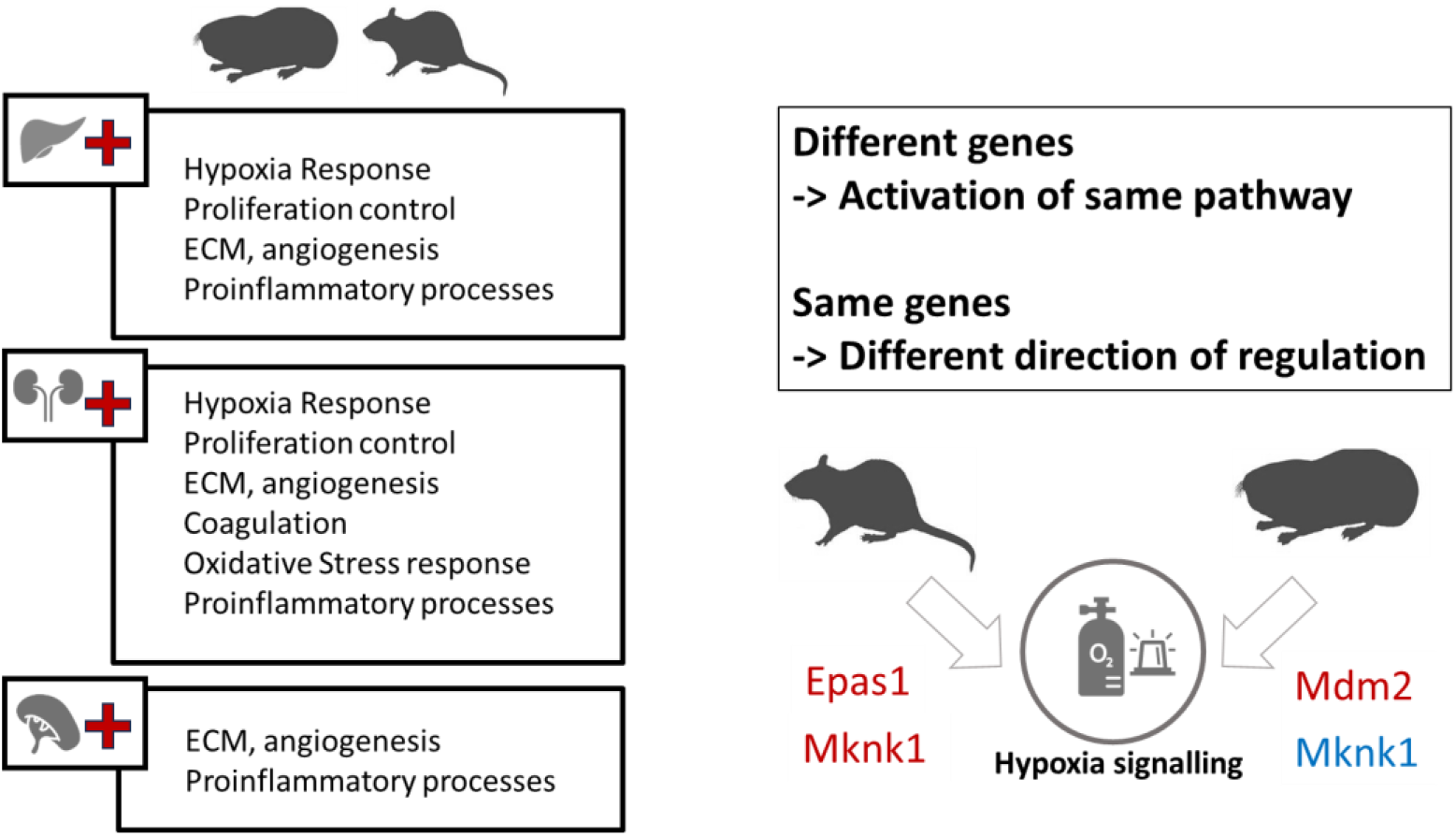
Similar biological pathways are activated in response to hypoxia in Spalax and rat organs (left). However, different genes may lead to activation of the same pathway (top right). Alternatively, the same gene can be regulated in different directions in response to hypoxia in the two species.

Differential gene expression under normoxic conditions is believed to be another important adaptive mechanism in *Spalax*. Elevated expression levels of important master regulators of hypoxic stress (Shams et al., 2004) and antioxidant defence (Schmidt et al., 2016) (Schülke et al., 2012) have been observed in the mole rat, suggesting that such higher expression levels of key genes and pathways, already present at normoxia, enable *Spalax* to react substantially faster towards an acute onset of severe hypoxic stress, which the rodents meet in their underground habitat during heavy seasonal rainfalls (Shams et al., 2005) (Schmidt et al., 2017). This can clearly be a useful adaptive strategy for biological processes like antioxidative defence or DNA-repair, which guarantee genomic integrity. For other biological processes such as inflammation or energy metabolism, a constitutive activation could be wasteful in energy expenditure and would not be in line with the energy-conserving phenotype of many subterranean species including *Spalax* (Lacey et al., 2001). It is therefore not surprising that we predicted an inhibition of biological pathways associated with inflammation, energy metabolism or angiogenesis in *Spalax* compared to rat (Tab. 4, Fig 5). These gene-regulatory patterns were in many cases observed in more than one organ, often utilizing the same pathway components as exemplified by the mTor signaling pathway (Tab. 5). This indicates the presence of body-wide adaptive gene-regulatory mechanisms acting in *Spalax*.

**Fig. 5:**
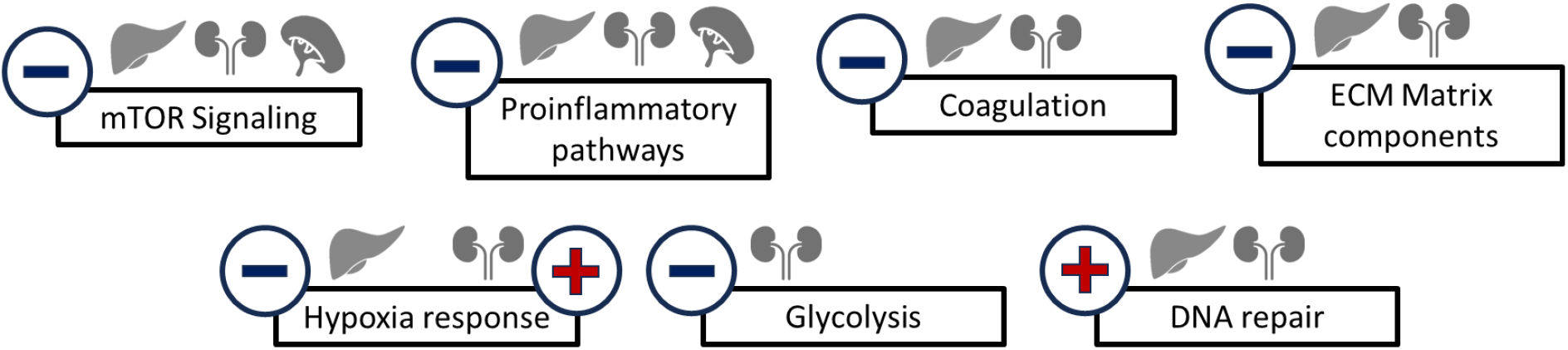
Predicted activation (red plus sign) or inhibition (blue minus sign) of pathways which are potentially associated with the Spalax phenotype at normoxic, unstressed conditions.

### Pathways involved in *Spalax* hypoxia adaptation: a detailed look

A more detailed look at the gene regulatory pathways involved in the differences between *Spalax* and rat revealed a number of observations that are of potential biological interest.

In both species and all three organs, we observed enrichment of terms associated with the *control of **cell death and proliferation***. This is an expected response to hypoxic stress in sensitive species, since hypoxia often leads to elimination of potentially damaged cells to prevent carcinogenesis (Lipton, 1999) (Yuan, 2009) (Coimbra-Costa et al., 2017). Hypoxia-adapted taxa, in contrast, might want to *avoid* unfavorable cell loss when encountering environmental stress repeatedly. In fact, a tight control of apoptosis has evolved in *Spalax* via mutations in the key tumour suppressor p53, some of which mimic mutations found in P53 of human tumours (Ashur-Fabian et al., 2004), or are convergently present in distantly related hypoxia-tolerant species (Domankevich et al., 2016). These changes reduce *Spalax* p53’s ability to induce apoptosis-regulating target genes like *Apaf1* and *Noxa* (Ashur-Fabian et al., 2004). In addition, *Mdm2*, a negative regulator of p53, was significantly higher expressed in *Spalax* than in rat under hypoxia in muscle, brain and heart tissue (Band et al., 2010). In our study, we observed a similar overexpression of *Mdm2* in *Spalax* liver and kidney, implying a body-wide involvement of this gene in controlling p53-mediated processes in the mole rat. While *Spalax* p53 appears to have a reduced stimulatory effect on pro-apoptotic genes, *in vitro* reporter gene assays revealed that it over-activates genes associated with cell-cycle arrest like *Cdkn1a* and *Pten,* suggesting an adaptive strategy that promotes temporary cell cycle arrest over apoptosis to prevent excess hypoxia-induced cell loss (Ashur-Fabian et al., 2004). Interestingly, we detected a lower transcription of *Pten* and *Cdkn1a* in all three *Spalax* organs, possibly down-modulating p53-induced activation under normoxic conditions. Thus, to improve insight into complex pathways, mutations in master regulators should be interpreted together with mRNA levels of their respective target genes. An additional factor that plays an important role in the cellular response to hypoxia is the calcium influx, which in turn, activates apoptosis and affects the transcription landscape. A study on *Spalax* cultured cells revealed a significantly reduced calcium influx during hypoxia (Salah-Hussiesy et al., 2018), implying a crucial role of maintaining homeostasis, rather than ‘overreacting’ under stress.

As expected, we observed an activation of classical hypoxia response terms like ***HIF1α signaling*** in liver and kidney in both species. Moreover, this pathway was predicted to be already activated under normoxic conditions in *Spalax* kidney. Adaptations in gene regulation in kidney have previously been proposed to play a role in *Spalax* hypoxia resistance, as the kidney is the major site of erythropoietin production. In this context, an elevated HIF1a and EPO production was measured in *Spalax* kidney (Shams et al., 2004), which likely contributes to the mole-rat’s elevated erythrocyte numbers (Arieli et al., 1986). Under hypoxia, non-adapted rodent species additionally show stress-erythropoiesis in the spleen (Wang et al., 2021), and the spleen serves as a reservoir of red blood cells in many diving species (Cabanac et al., 1997). Interestingly, *Spalax* spleen showed a lower transcriptional response to hypoxia and fewer interspecies differences in pathway activation under normoxic condition, as compared to the other organs. This hints at an obsolescence of splenic stress erythropoiesis in *Spalax* due to its well-adapted erythropoietic physiology.

An important cause of cell and DNA damage is **oxidative stress,** which arises from cycles of acute hypoxia followed by re-oxygenation and produces harmful reactive oxygen species (ROS) (Guzy & Schumacker, 2006). Hypoxia induced upregulation of enzymes connected with defence against oxidative stress, like *Hmox1*, can therefore be observed in many species in response to hypoxia (Panchenko et al., 2000). Our RNA-Seq and qPCR data confirmed an elevated mRNA level of *Hmox1* in *Spalax* and rat in response to hypoxia, indicating that despite the many adaptations in gene regulation under oxygen deprivation, common conserved regulatory mechanisms can be found. While basal levels of oxidative stress can be found in essentially all aerobic organisms, they are substantially increased in subterranean, diving and high-altitude species (Eaton & Pamenter, 2022). *Spalax* faces acute cyclical changes in oxygen levels in its habitat when performing energy-consuming digging activities in confined tunnels at one moment and entering better-ventilated burrow areas at the next. Additionally, the mole rat meets an acute severe hypoxic condition when its burrow systems are flooded. As a potentially adaptive difference from the rat, we detected overexpression in *Spalax* of *Hmox1* in kidney tissue, which had also been previously observed in liver, heart and brain tissue ((Schülke et al., 2012; Tab S9 present study).

ROS that are not sufficiently buffered by the radical scavenging system represent a severe threat to genomic integrity, producing various types of DNA damage like nucleotide modifications or crosslinks (Jena, 2012) (Renaudin, 2021) (Habibi et al., 2022). Furthermore, severe hypoxia induces S-phase arrest and dNTP depletion, leading to additional replicative stress (Pires et al., 2010) (Gelot et al., 2015) (Ramachandran et al., 2021). Consistent with this, we observed an enrichment of ***DNA damage response-associated processes*** among the genes that were differentially expressed at normoxia between the two species. Of note, “Base excision repair” and “Role of BRCA1 in DNA damage response” were predicted as activated in liver and kidney of the mole rat. Previous analyses of RNA-Seq data from *Spalax* liver (Schmidt et al., 2017) and brain (Malik et al., 2016) had indicated an elevated expression in the mole rat of key genes from the Fanconi Anemia pathway, which orchestrates different DNA repair activities (Peake & Noguchi, 2022). In the present study, we were able to confirm these results for kidney and spleen as well (Fig. 6), indicating an organ-overarching adaptation of DNA repair processes in *Spalax*.

**Fig. 6:**
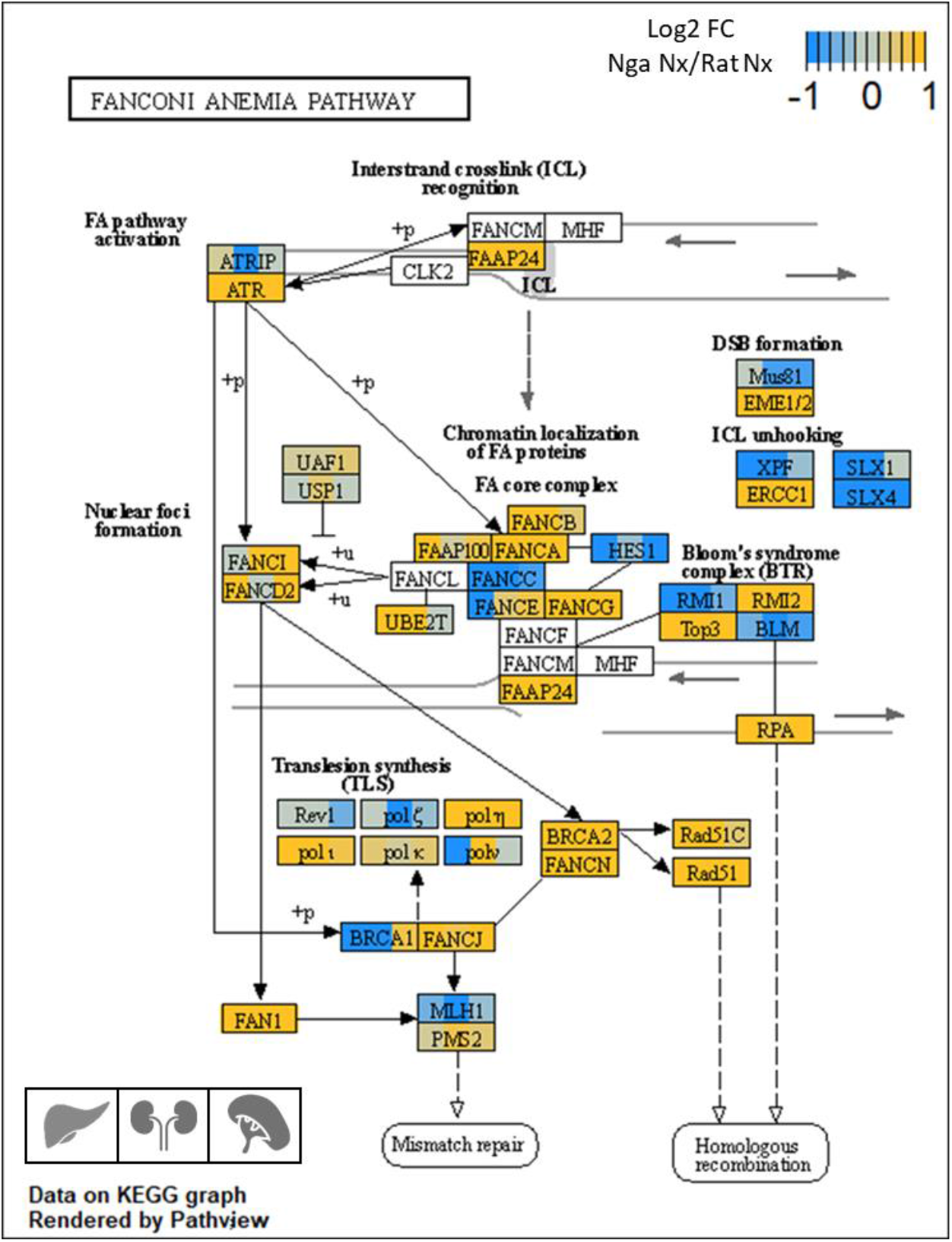
Differential expression between Spalax (Nga) and rat (Rno) of genes contributing to the Fanconi anemia pathway in liver, kidney and spleen (respective organs are represented from left to right in the boxes). Genes with higher expression in Spalax are overrepresented compared to genes with a lower expression. Differential expression is represented in form of the Log2 FC of expression levels between Spalax under normoxia and rat under normoxia (Log2 FC (Nga Nx/Rno Nx)). Genes with higher expression in Spalax are marked orange.

Looking for candidate genes of special biological relevance in these pathways, important DNA repair-associated genes like the master activator *Ataxia telangiectasia and Rad3 related (Atr),* the *breast cancer type 2 susceptibility protein (Brca2)* and the *Werner syndrome ATP dependent Recql1 helicase* (*Wrn*) were found overexpressed in all three analysed organs in *Spalax* compared to rat (present study; (Schmidt et al., 2017) as well as in brain tissue (Malik et al., 2016), suggesting body-wide elevated activities of their gene products. Recently, DNA damage repair assays performed on skin fibroblasts isolated from *S. carmeli* (another subspecies of the *S. ehrenbergi* taxon) indeed revealed an elevated resistance against H_2_O_2_, the topoisomerase inhibitor etoposide and UV-C radiation-induced damage compared to rat fibroblasts (Domankevich et al., 2018), suggesting an accelerated repair of those lesions in S*palax*. Since DNA damage is causal to both cancer development (Torgovnick & Schumacher, 2015) and ageing (Schumacher et al., 2021), we hypothesize that an improved capacity of recovering DNA lesions may substantially contribute to *Spalax’s* anti-cancer phenotype and its longevity.

An important physiological adaptation to its hypoxic habitat is *Spalax’s* low basal metabolic rate, which reduces its oxygen consumption and–thereby–oxidative stress (Widmer et al., 1997). Such hypometabolism, which is thought to promote longevity and prevent cancer (Garasto et al., 2017), was in fact previously inferred from transcriptome analyses of *Spalax* brain, where lower mRNA levels of genes associated with energy metabolism were found compared to rat (Malik et al., 2016). In support of this, we detected differential expression of genes associated with cellular ***energy metabolism*** in liver, kidney and spleen between *Spalax* and rat at normoxia. For example, genes contributing to the oxidative respiration (Fig.7) like *cytochrome c oxidase*, the key enzyme complex responsible for oxygen consumption, showed lower normoxic mRNA expression for most subunits in all analysed *Spalax* organs.

**Fig. 7:**
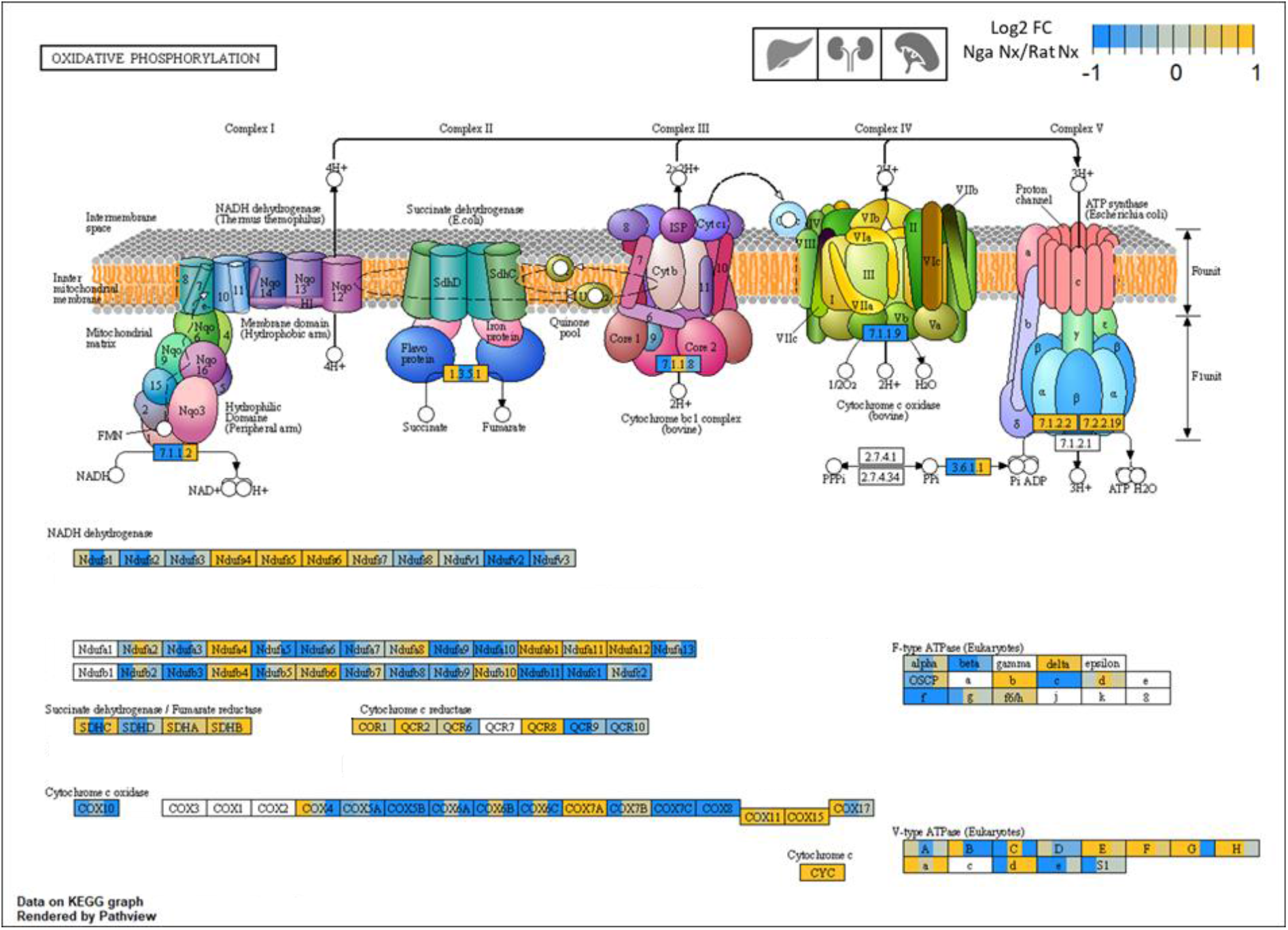
Differential expression between Spalax (Nga) and rat (Rno) of genes contributing to the oxidative phosphorylation in liver, kidney and spleen (respective organs are represented from left to right in the boxes). Genes with lower expression in Spalax are overrepresented compared to genes with a higher expression. Differential expression is represented in form of the Log2 FC of expression levels between Spalax under normoxia and rat under normoxia (Log2 FC (Nga Nx/Rno Nx)). Genes with higher expression in Spalax are marked orange.

Another potential way of hypoxia adaptation is to increase oxygen utilization by ***modification of catabolic pathways***, e.g., by switching off the energy substrate. Under oxygen deprivation, naked mole rats utilize anaerobic metabolism fueled by fructose (Park et al., 2017). In *Spalax*, glucose metabolism is likely switched already under normoxic conditions from oxidative pathways to pentose phosphate pathways, lactate production and cataplerotic pathways (Miskevich et al., 2021). In our present study, we observed that many key genes for anaerobic metabolism like triose-phosphate isomerase (TIM), pyruvate kinase M (PKM), alcohol dehydrogenase 5 (ADH5) and lactate dehydrogenase B (Ldhb) were overexpressed in all three analysed *Spalax* organs. In brain, genes associated with glycolysis, like the key enzyme hexokinase, were downregulated (Malik et al., 2016). In this context, our transcriptome data predicted an inactivation of “Glycolysis” and “Glycogen degradation*”* in *Spalax* kidney under normoxia, indicating further adaptation of glycogen metabolism in the mole rat. In liver, however, we predicted a constitutive activation of glycogen degradation. In this context, Nasser et al., (2009) detected especially high glucose level in the serum of *Spalax* compared to human blood. The authors discussed these findings as a potential result of insufficient fasting of the animals before blood collection. With respect to the results of the present study, the elevated blood glucose level might be due to constitutive overexpression of genes associated with elevation of blood glucose, which could be adaptive to *Spalax* in its hypoxic environment by providing higher concentration of substrate for energy metabolism.

Recent studies on the hypoxia-adapted rodent Qinghai voles (*Neodon fuscus*) suggested an improvement of **fatty acid-based oxidative phosphorylation** in this species after hypoxia exposure (Li et al., 2021). As a potential parallel, in *Spalax* kidney we predicted a constitutive activation of pathways associated with fatty acid oxidation, possibly indicating another way of metabolic adaptation in the mole rat.

Importantly, the transcriptome data also predicted an inactivation of **mTor signaling** in all three organs in *Spalax* compared to rat. Due to its involvement in ageing-related processes like nutrient sensing, proteostasis and mitochondrial dysfunction, the mTOR cascade is considered a major player in life span regulation (Papadopoli et al., 2019) (Weichhart, 2018) (Bjedov & Rallis, 2020) and mTOR inhibitors like rapamycin are researched for their potential use in lifespan extension (Blagosklonny, 2019). Interestingly, recent studies in other hypoxia-adapted species (naked mole rat and red-eared slider turtles (Wu & Storey, 2021)) suggested an *activation* of mTOR in response to hypoxia, which may aid metabolic reprogramming in favor of anaerobic metabolism. In *Spalax,* specific amino-acid replacements in mTORs functional domains were observed (Schmidt et al., 2017), suggesting potential adaptations on the functional protein level in the blind mole rat. A constitutive inactivation of the mTOR pathway, as inferred from the transcriptome level in *Spalax*, could mimic the effect of anti-cancer drugs (Hua et al., 2019) (Bjedov & Rallis, 2020) and thus contribute to the mole rat’s cancer-resistance.

Hypoxia plays a key role in the regulation of **immunity and inflammation** since it is a hallmark of inflamed, infected or damaged tissue (Villafuerte et al., 2021) . In this context, inflammation can induce the activity of hypoxia-response pathways, and hypoxia may modulate inflammatory signaling (Wanderer, 2011). It is therefore not surprising that we detected an enrichment of genes associated with the regulation of inflammatory processes among the hypoxia-responsive genes in all organs of both species. The same was also observed in brain tissue in previous studies (Malik et al., 2012). Sterile inflammation, which is induced by the secretome of senescent cells consisting of inflammatory cytokines, chemokines and growth factors, knowingly plays a role in age-related disorders and cancer (Franceschi et al., 2018) (Teissier et al., 2022). Stress assays in *Spalax* fibroblasts suggest, that the positive feedback loop of IL1α-NF-κB, which is an upstream regulatory machinery of the inflammatory response, appears to be impaired (Odeh et al., 2020). Accordingly, we detected in all analysed *Spalax* organs a decreased expression of proinflammatory factors, some of which are also part of the secretome of senescent cells (Il1a, Il6r and Cxcl1 (Saul et al., 2022). We conclude that a lower expression of proinflammatory genes under normoxic conditions in healthy organs may produce an “inflammation-free” phenotype in *Spalax*, possibly contributing to the species’ longevity.

**Angiogenesis** is an important mechanism to ensure adequate oxygen supply (Griffioen & Dudley, 2021). Not unexpectedly, we found that hypoxia-responsive differentially expressed genes were associated with angiogenesis in all three organs in both species. Elevated *Vegfa* expression was suggested to improve vascularization in *Spalax* muscle tissue, thereby ameliorating oxygen transport (Avivi et al., 2005). In the present study, we observed a similar constitutive, normoxic overexpression of *Vegfa* in *Spalax* kidney and spleen. Angiogenesis is tightly regulated by a variety of antiangiogenic factors such *Calcipressin1* (Rcan1) (Huang & Bao, 2004) (Baek et al., 2009). Previous studies observed a 10-fold upregulation of Rcan1 mRNA-expression in *Spalax* muscle, but not in rat muscle (Malik et al., 2012), which we also detected in liver and kidney transcriptomes. In addition, our data predicted an inactivation of pathways associated with extracellular matrix degradation (“Chondroitin- and Dermatan Sulfate Degradation”) under normoxic conditions in liver and kidney. This is in line with results of *in vitro* assays suggesting that alternatively spliced heparanases might contribute to an inhibited ECM degradation in *Spalax* (Nasser et al., 2005) (Nasser et al., 2009). Taken together, these observations suggest a sophisticatedly evolved regulation of angiogenesis mechanisms to create an extensively vascularized tissue, while concurrently inhibiting a pro-metastatic environment (Malik et al., 2012) (present study).

### Convergent patterns of gene expression indicate adaptive processes

As an important caveat to our (and other) transcriptomic interspecies comparisons, it might be argued that the observed differences between taxa simply reflect phylogenetic distance, but not necessarily adaptive processes. A way out of this dilemma can be the inclusion of additional species with and without the studied phenotypes, which may reveal instances of phenotypic convergence, and thus indirectly point at ‘adaptation in action’.

In rare cases, convergent adaptation is realized by ‘strict convergency’, e.g., by mutations in *the same* gene in two species adapted to the same selective pressure. One example in *Spalax* is the presence of functionally similar amino acid replacements in the proton-gated nociceptor sodium channel Nav1.7 (Scn9a) in several species adapted to hypercapnic environments (Fang et al., 2014). More often, convergency is realized on a functional level by sometimes utilizing different genes to achieve similar phenotypes. In the context of hypoxia, adapted species like naked mole rats or whales (Toren et al., 2020) show differential expression in genes associated with processes like oxidative response, metabolism, ROS defence and DNA-repair compared to non-adapted related taxa (Yu et al., 2011) (Lewis et al., 2013) (MacRae et al., 2015) (Heinze et al., 2018). Since we observed gene-regulatory changes influencing the very same biological processes in *Spalax*, we are confident that these differences in gene expression are to a high degree adaptive. There are even examples for direct convergence in overexpression of the same genes in *Spalax* and naked mole rat, as exemplified by alpha2-macroglobulin A2M (Schmidt et al., 2017) (Thieme et al., 2015) and DNA-repair genes like ATR (MacRae et al., 2015). Studying and comparing functional and direct convergencies in adapted species like *Spalax* and naked-mole rat can thus provide valuable insight into mechanisms of evolution and help to pinpoint different ways to modulate physiological processes like hypoxia response or tumour inhibition that are of high biomedical interest.

## Conclusion

*Spalax* adaptations to its challenging subterranean habitat are likely influenced by transcriptomic changes, acting overarchingly across vital organs and tissues. These transcriptomic changes involve genes from fundamental biological processes like the control of cell death, ROS defence, DNA repair, energy metabolism, immune response and angiogenesis. Many of these processes play a role in hypoxia response, but are knowingly also involved in the defence against cancer, the maintenance of genomic integrity and the emergence of longevity. Our data thus provide additional evidence for the hypothesis that these biomedically most interesting phenotypes have evolved as byproducts of *Spalax’* hypoxia adaptation. Future studies on cis- and trans regulatory mechanisms orchestrating the adaptation on the gene regulatory level in *Spalax* will provide further insights in the evolution of the mole rat’s remarkable complex phenotype.

## Acknowledgements

We would like to thank Alexandra Hassemer and Simon Steines for contributing to the qPCR-analyses. This project was funded by the Deutsche Forschungsgemeinschaft (DFG, German Research Foundation)—GRK2526/1—Project no. 407023052.

