## Supplementary data 2 for "Adaptation of the *Spalax galili* transcriptome to life under hypoxia may hold a key to a complex phenotype including longevity and cancer resistance"

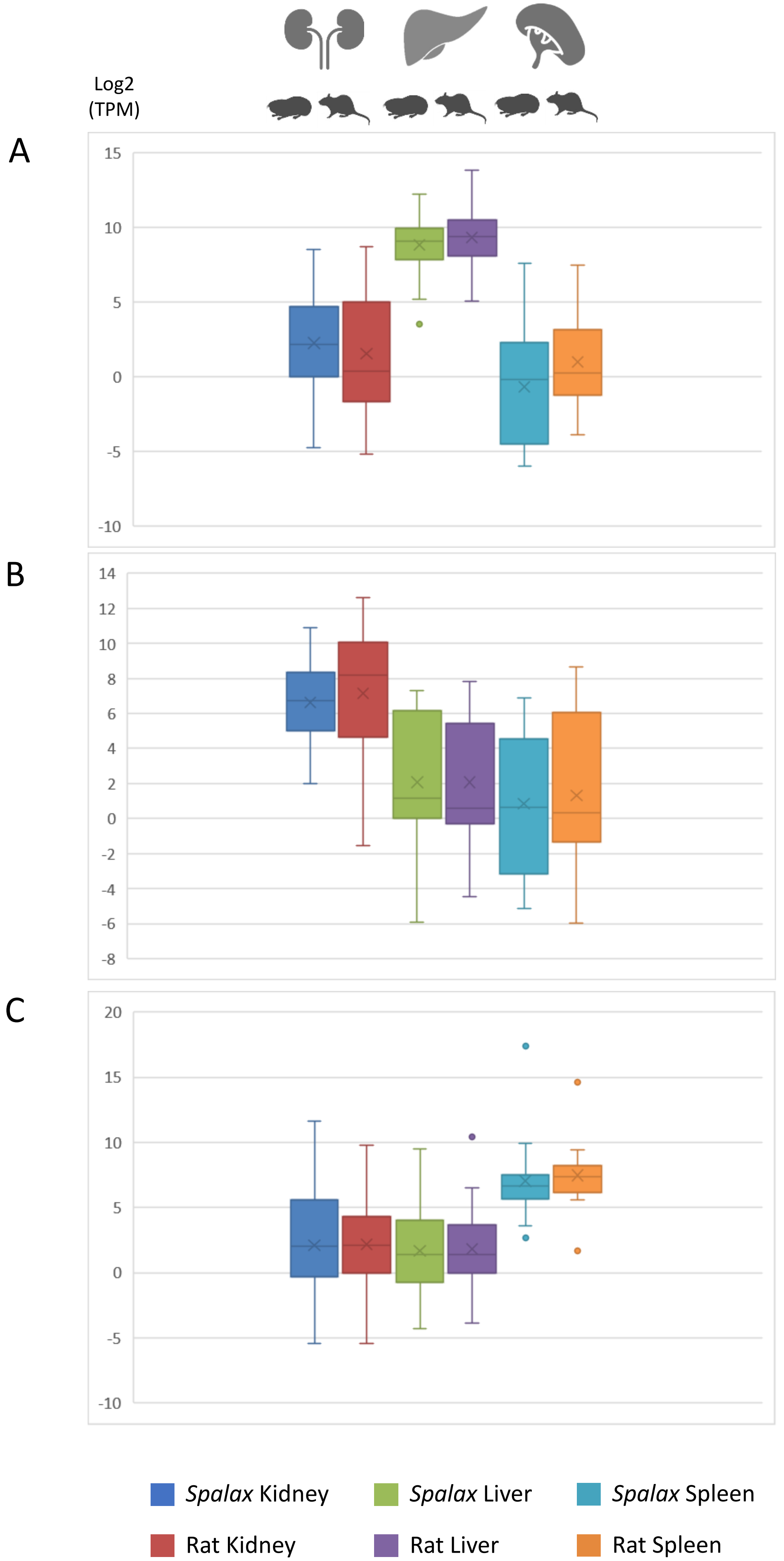
Log2
(TPM)
A
15
10
5
0
-5
-10
B
14
12
10
8
6
4
2
0
-2
-4
-6
-8
C
20
15
10
5
0
-5
-10
Spalax Kidney
Rat Kidney
Spalax Liver
Rat Liver
Spalax Spleen
Rat Spleen

Expression of organ specific genes (Yu et al., 2014) in *Spalax* and rat. A = Liver-specifically expressed genes, B = Kidney-specifically expressed genes, C = Spleen-specifically expressed genes
