## Supplementary data 4 for "Adaptation of the *Spalax galili* transcriptome to life under hypoxia may hold a key to a complex phenotype including longevity and cancer resistance"

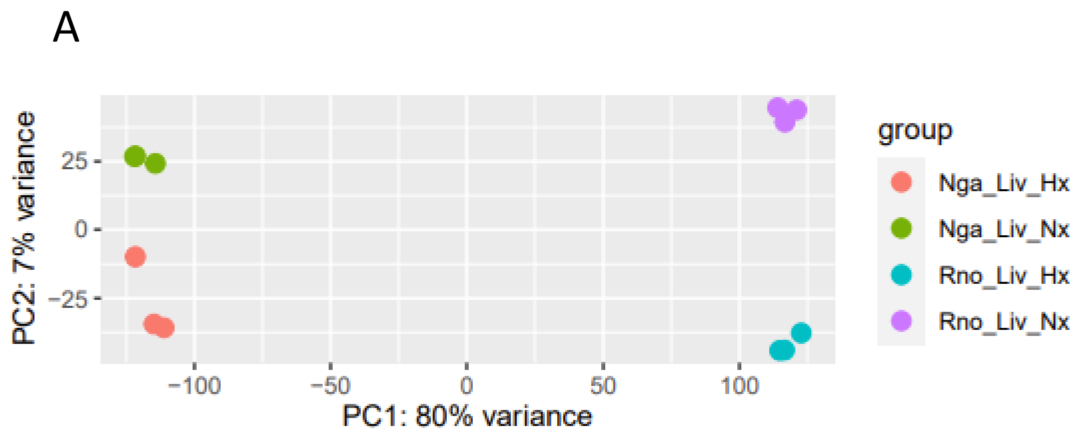


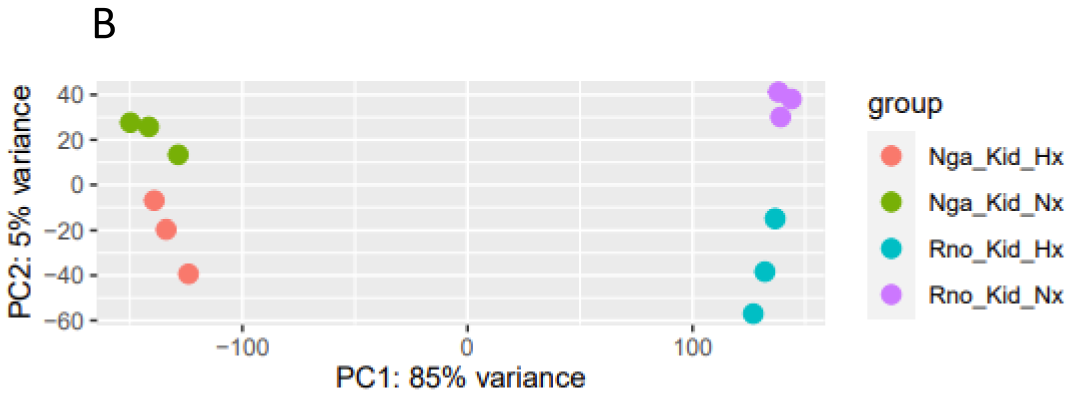


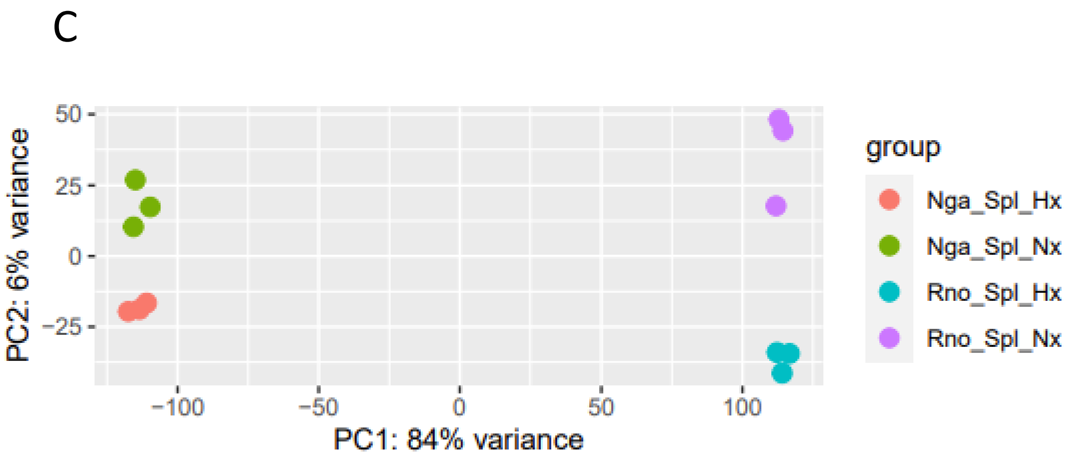


Principal component analyses of trancriptome data of *Spalax* (Nga) and Rat (Rno) under normoxic- (Nx) and hypoxic (Hx) conditions in A = Liver (liv), B = Kidney (Kid) and C = Spleen (Spl)
