## Supplementary data 5 for "Adaptation of the *Spalax galili* transcriptome to life under hypoxia may hold a key to a complex phenotype including longevity and cancer resistance"

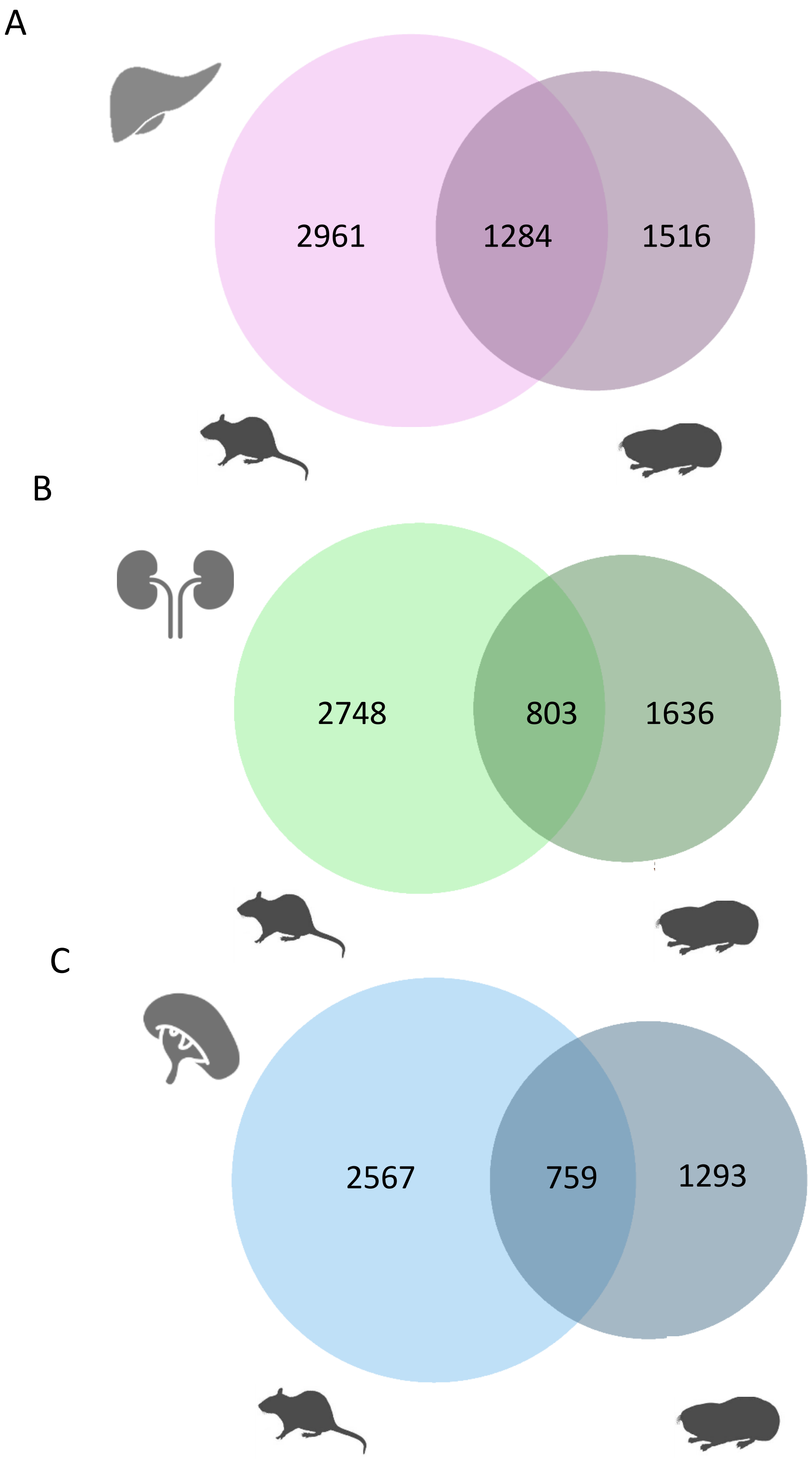


*Hypoxia regulation of gene expression compared through all organs (padj < 0.05). Genes differentially expressed under hypoxia in Spalax and rat in A liver, B kidney, C Spleen*
