## Supplementary data 8 for "Adaptation of the *Spalax galili* transcriptome to life under hypoxia may hold a key to a complex phenotype including longevity and cancer resistance"

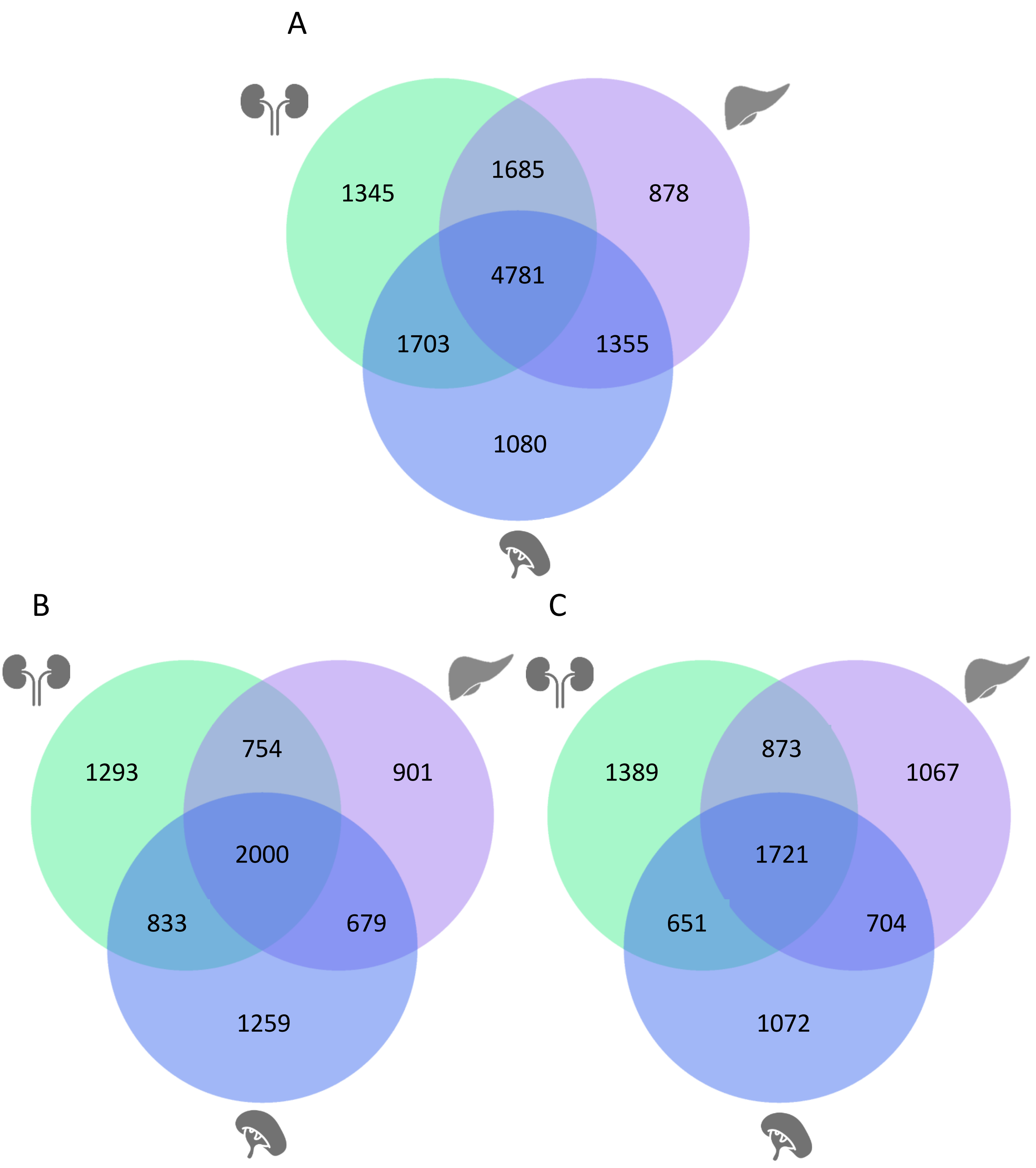


*Differential gene expression in Spalax versus rat under normoxia in all organs (padj < 0.05). A = All differentially expressed genes, B = Genes higher expressed in Spalax compared to rat, C = Genes lower expressed in Spalax compared to rat*
