## Supplementary data 9 for "Adaptation of the *Spalax galili* transcriptome to life under hypoxia may hold a key to a complex phenotype including longevity and cancer resistance"

Relative expression of selected candidate genes between hypoxic (Hx) and normoxic (Nx) *Spalax galili* and *Rattus norvegicus* kidney, spleen and liver samples quantified by qRT-PCR and RNA-Seq. Green shading indicates matching results, grey shading indicates diverging results, *= padj. < 0.05, # = padj. > 0.05 low = high ratio due to low copy number at normoxia,

| **Kidney** | **Spalax Hx/Nx** | | **Rat Hx/Nx** | | **Spalax Nx/ Rat Nx** | |
|---|---|---|---|---|---|---|
| **Gene** | qRT-PCR | RNA-Seq | qRT-PCR | RNA-Seq | qRT-PCR | RNA-Seq |
| Gpnmb | 1.13 | 2.04* | 0.82 | 2.12* | 2.63 | 3.92* |
| Fen1 | 1.36 | 0.34 | 0.53 | 0.17 | 1.78 | 3.1* |
| Wrn | -0.59 | -0.42 | 0.85 | 0.34 | 3.94 | 1.73* |
| Hmox1 | 4.01 | 4.7* | 3.17 | 4.12* | 7.28 | 3.15* |

| **Spleen** | **Spalax Hx/Nx** | | **Rat Hx/Nx** | | **Spalax Nx/ Rat Nx** | |
|---|---|---|---|---|---|---|
| **Gene** | qRT-PCR | RNA-Seq | qRT-PCR | RNA-Seq | qRT-PCR | RNA-Seq |
| Vegfa | 0.04 | -0.11 # | 2.68 | 2.46* | 2.16 | 1.47* |
| Fen1 | -3.54 | -0.01 | -0.11 | -0.02 | 3.04 | 1.6* |
| Wrn | -2.3 | -0.45 | -0.45 | -0.43 | 4.13 | 2.44* |
| Pnkp | -0.86 | -0.16 | -0.63 | -0.64* | 2.72 | 0.71* |

| **Liver** | **Spalax Hx/Nx** | | **Rat Hx/Nx** | | **Spalax Nx/ Rat Nx** | |
|---|---|---|---|---|---|---|
| **Gene** | qRT-PCR | RNA-Seq | qRT-PCR | RNA-Seq | qRT-PCR | RNA-Seq |
| A2m | 0.49 | 0.74 | 6.56 | 3.9* | 10.57 | 15.37* |
| Atr | 0.26 | -0.06 # | 0.58 | 0.65* | 4.84 | 2.89* |
| Cisd2 | 0.38 | -0.14 # | 0.77 | 0.56 | 5.28 | 2.05* |
| Fgf21 | 8.18 | 7.13* | 0.77 | -0.54 # | 0.14 | -5.47*low |
| Wrn | 0.26 | 0.02 | -1.00 | -0.37 | 5.30 | 2.39* |
| Xpa | -0.15 | -0.45 | -0.74 | -0.70* | 4.63 | 3.33* |
| Rcan1 | 3.41 | 2.12* | -1.32 | -1.70* | -0.22 | -1.24* |
